# Scaffolding of RhoA contractile signaling by anillin: a regulatory analogue of kinetic proofreading

**DOI:** 10.1101/282756

**Authors:** Srikanth Budnar, Kabir B. Husain, Guillermo A. Gomez, Maedeh Naghibosidat, Suzie Verma, Nicholas A. Hamilton, Richard G. Morris, Alpha S. Yap

## Abstract

Scaffolding is a fundamental principle of cell signaling commonly thought to involve multi-domain proteins that tether different components of a pathway together into a complex ^1,2^. We now report an alternative mechanism for scaffolding that is necessary for RhoA-mediated contractile signaling. We find that anillin binding stabilizes active, GTP-RhoA, and promotes contractility at both the epithelial zonula adherens (ZA) and the cytokinetic furrow. However, anillin does not conform to the classical picture of a multi-domain tether, since its RhoA-binding AH domain alone was sufficient to promote contractile signaling. Moreover, anillin competes with contractile effectors for a common site on RhoA, presenting the conundrum of how an inhibitory interaction can otherwise promote signaling. To explain this, we propose that inactivation of RhoA is non-Poissonian, having a rate that increases with time, unless the process is reset via transient binding to anillin. Repeated cycles of binding and un-binding therefore increase cortical residence times of non-sequestered GTP-RhoA and hence the probability of engaging contractile effectors. We identify the modification of the local lipid environment as a potential mechanism underlying such non-Poisson statistics, and demonstrate agreement with a minimal cellular system. Finally, we show that Myosin II anchors anillin at the cortex to form a feedback pathway that enhances RhoA signaling. This new paradigm of scaffolding is a regulatory analogue of kinetic proofreading and may be employed by other binding proteins that do not fit the classical picture.

---

Cellular contractility requires strict spatio-temporal control of signaling within cells. Contractile zones such as the cytokinetic furrow and zonula adherens (ZA) of epithelial cells are controlled by localized RhoA, which, in its active, GTP-loaded form, recruits multiple effector proteins to assemble and maintain the actomyosin apparatus at the cell cortex ^3^. In turn, GTP-RhoA is classically thought to be controlled by molecules that regulate its generation, inactivation, or dissociation from the membrane. Anillin is a putative scaffold, first identified at the contractile furrow during cytokinesis ^4,5^ and increasingly implicated in contractility elsewhere, such as at cell-cell junctions in Xenopus embryos ^6^ and as a component of the E-cadherin interactome ^7^. Anillin can interact with a diverse range of proteins, notably directly with GTP-RhoA, F-actin, and non-muscle Myosin II (NMII) ^8–10^. Thus, anillin has been thought to function as a RhoA effector, being recruited to the cortex by active RhoA, then regulating contractility by binding F-actin and NMII ^5,9^.

Anillin preferentially localized with actomyosin at the ZA of confluent MCF-7 epithelial cells (Fig. 1a; Extended data Fig. 1a) and junctional tension, measured by recoil following laser ablation, was reduced by anillin shRNA (knock-down, KD) (Fig. 1b,c; Extended Data Fig. 1b). This suggested that anillin contributes to junctional contractility, a conclusion that was reinforced by reduced levels of actomyosin and the RhoA effectors, ROCK1 and mDia1, at anillin KD junctions (Extended Data Fig. 1c, e-i). The specificity of anillin KD was confirmed by reconstitution with an RNAi-resistant WT anillin transgene (anillin^WT^). Interestingly, junctional NMIIA was also recovered in KD cells expressing an anillin mutant lacking the actin-binding domain, but not when either the NMII-binding domain (anillin^ΔMyo^) or the AH domain (anillin^ΔAH^) were deleted (Fig. 1f; Extended Data Fig. 2 a,f). However, when corrected for junctional levels of the transgenes, anillin^ΔMyo^ restored junctional NMIIA as effectively as anillin^WT^, but a defect persisted with the ΔAH mutant (Extended Data Fig. 2 g). This suggested that the NMII-binding domain might function by localizing anillin while the AH domain supported another mechanism.

**Figure 1.**
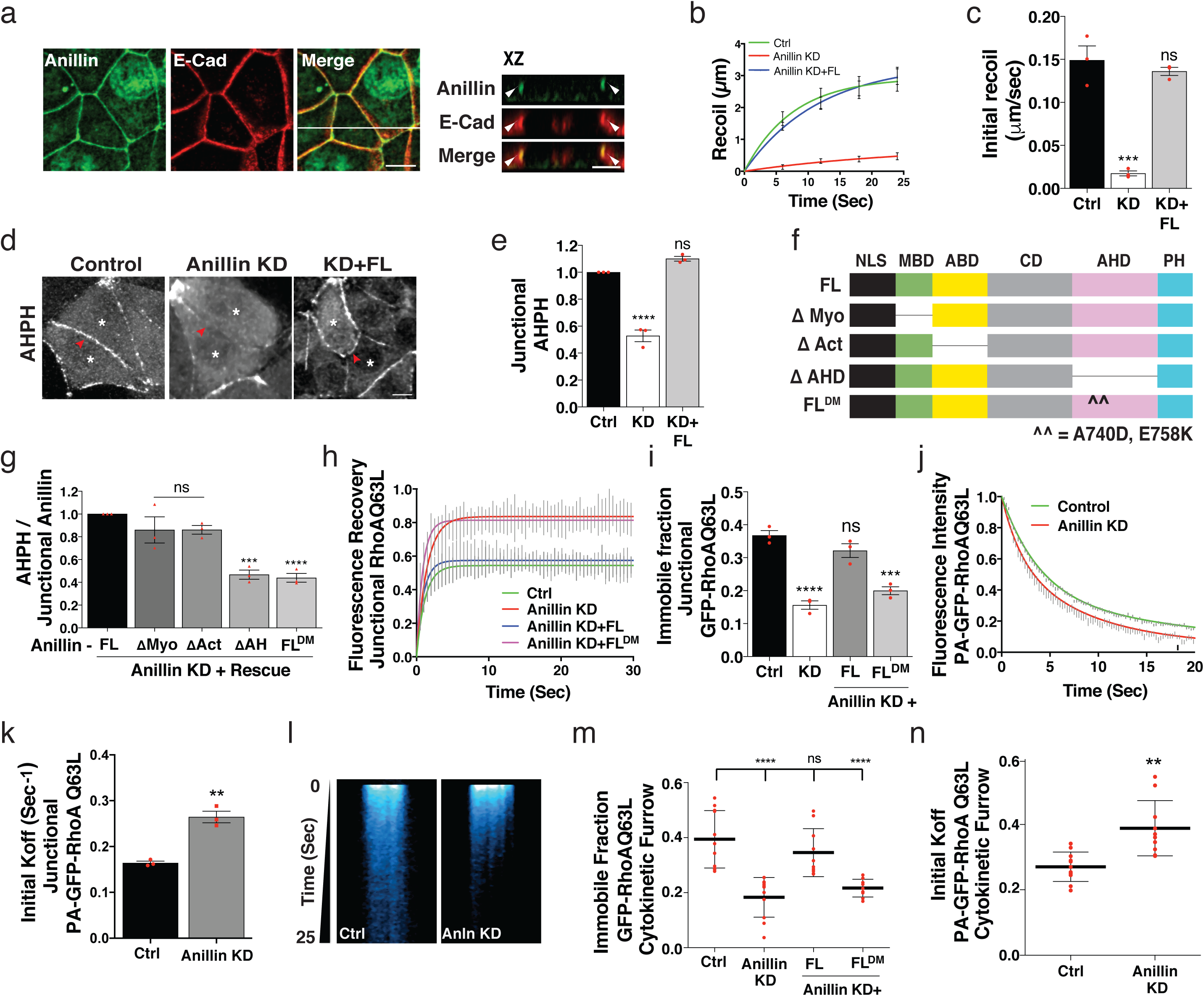
Anillin binds and stabilizes active RhoA at sites of contractility. (**a**) Co-localization of endogenous anillin with E-Cadherin; XZ views, taken at the indicated lines, show apical accumulation of anillin with E-cadherin. (**b,c**) Recoil measurements (**b**) and initial recoil velocity (**c**) of cell junctions after laser ablation of junctions. (**d,e**) Immunostaining (**d**) and fluorescence intensity (**e**) of junctional AHPH in MCF-7 cells expressing control shRNA, Anillin shRNA (KD), or Anillin shRNA reconstituted with full-length RNAi resistant Cherry-anillin (Anillin^WT^). (**f**) Domain structure of full-length (FL) anillin and the various mutant transgenes used in this study. (**g**) Fluorescence intensity of GFP-AHPH at ZA in anillin shRNA MCF-7 cells normalized to the expression levels of the reconstituted anillin transgenes. FL^DM^ = FL^A740D,E758K^ (**h,i**) FRAP of GFP-RhoAQ63L at the ZA in Anillin KD or reconstituted cells: best-fit curves (**h**), immobile fractions (**i**). (**j-l**) Fluorescence decay (**j**), initial Koff (**k**), and kymograph (**l**) of photoactivated RhoAQ63L (PA-RhoAQ63L) at ZA in control or Anillin shRNA MCF-7 cells. (**m**) FRAP of GFP-RhoAQ63L at the cytokinetic furrow in Anillin KD or reconstituted cells; immobile fractions from best-fit curves. (**n**) Initial Koff of of photoactivated RhoAQ63L (PA-RhoAQ63L) at the cytokinetic furrow in control or Anillin shRNA MCF-7 cells. Data represent means ± s.e.m and n = 3 independent experiments except for (**m**) and (**n**) where data represents ± s.d and n ≤11 cells. ** P < 0.01, ***P < 0.001, ****P < 0.0001; ns, not significant; One-way ANOVA with Dunnett’s multiple comparisons test (**c,e,g,i,m**); Students *t*-test (**k,n**). Scale bars, 10 μm; 5 μm for XZ views.

**Figure 2.**
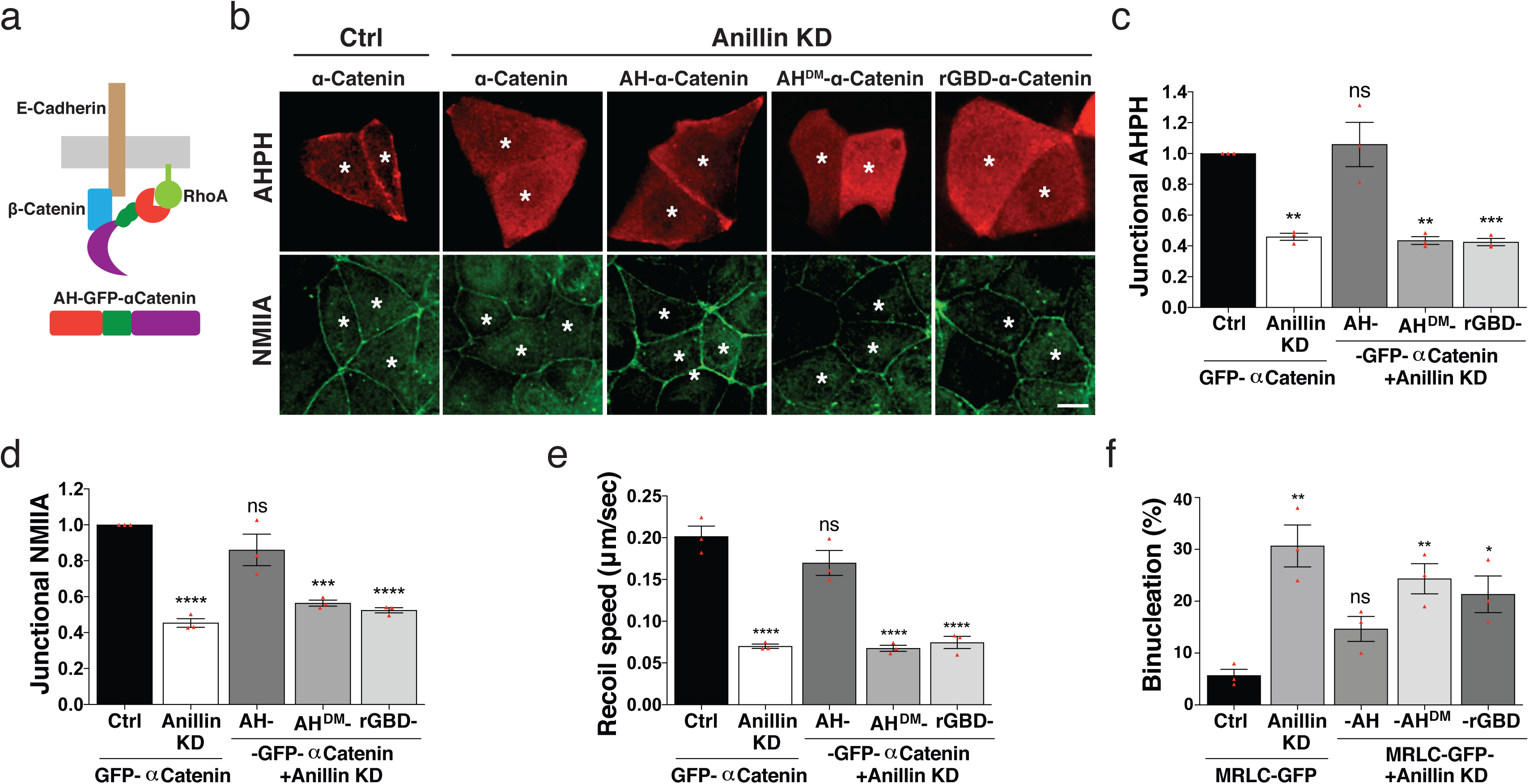
The anillin AH domain is sufficient to support junctional contractility. (**a**) Cartoon depicting the GFP-AH-α-catenin chimeric construct. (**b,c**) Representative images (**b**) and fluorescence intensity of GFP-AHPH (**c**) and NMIIA (**d**) at ZA of cells expressing control siRNA, Anillin siRNA (KD) and Anillin siRNA along with the indicated α-catenin chimeras (AH, AH^DM^ and rGBD). Asterisks indicate cells expressing the transgene. (**e**) Junctional tension (initial recoil velocity) in cells expressing the indicated siRNA and transgenes. (**f**) Quantification of cytokinetic defects (bi-nucleation) in MCF-7 cells expressing control siRNA, anillin siRNA (KD) and anillin siRNA along with the indicated MRLC chimeras (- AH, -AH^DM^ and -rGBD). Data represent means ± s.e.m and n = 3 independent. *P < 0.05, ** P < 0.01, ***P < 0.001, ****P < 0.0001; ns, not significant; One-way ANOVA with Dunnett’s multiple comparisons test. Scale bars, 10 μm.

One possibility was that the AH domain regulated RhoA itself, which is essential for junctional contractility ^11^. We tested this using AHPH, a location biosensor for GTP-RhoA ^9,12^ that derives from the C-terminus of anillin, but does not compete with anillin^WT^ for junctional localization (Extended Data Fig. 2 h,i). Anillin co-accumulated with AHPH and TCA-resistant RhoA staining at the ZA (Extended Data Fig. 1a). However, both markers of junctional RhoA were reduced ∼ 50% in anillin KD cells, but restored with anillin^WT^ (Fig. 1 d,e; Extended Data Fig. 1c,d). We confirmed that anillin was required for junctional RhoA, by developing two additional location biosensors based on the GTP-RhoA binding domains (GBDs) of mDia1 (mDia1-GBD) and ROCK1 (ROCK1-GBD). Both sensors localized to the ZA in a RhoA-sensitive fashion and were reduced by anillin KD (Extended Data Fig. 2 j-n). Strikingly, although junctional RhoA was reduced in cells expressing anillin^ΔMyo^ or anillin^ΔAH^, only the ΔAH mutant had a persistent effect after correction for junctional recruitment (Fig. 1g; Extended Fig. 3 a,b). This strongly implied that the anillin AH domain supports junctional contractility via RhoA.

Although the AH domain can interact with Ect2 ^5^, which activates RhoA at the ZA^11^, junctional Ect2 was not altered by anillin KD (Extended Data Fig. 3 c,d). Nor did anillin KD affect output from a FRET-based RhoA activity sensor (Extended Data Fig. 3 e), as might have been expected if anillin were regulating the balance of GEFs and GAPs that acted upon junctional RhoA. We therefore hypothesized that direct binding by its AH domain might allow anillin to stabilize GTP-RhoA at the junctional membrane. Indeed, FRAP experiments showed that anillin KD destabilized GFP-RhoA at the ZA, as well as RhoA^Q63L^, a GTPase-defective mutant (Extended Data Fig. 3 f,g; Fig. 1h,i), which excluded the possibility that anillin was regulating RhoA dynamics indirectly through changes in the kinetics of its activation and inactivation ^13^. This was confirmed by analysis of fluorescence decay after photoactivating RhoA^Q63L^ tagged with photoactivatable (PA)-GFP (Fig. 1j; Extended Data Fig. 3h). Anillin KD increased the apparent dissociation rate (Koff) of PA-GFP-RhoA^Q63L^, calculated from the initial rate of fluorescence decay (Fig. 1k). Kymographs showed little lateral diffusion either in control or anillin KD cells (Fig. 1l), implying that fluorescence decay principally reflected the dissociation of GTP-RhoA from the plasma membrane.

Together, these findings suggested that anillin might regulate RhoA at the ZA by direct binding, rather than through the canonical regulation of its nucleotide-bound status^14^. Indeed, mutating two residues in the AH domain (A740D/E758K) that are required for GTP-RhoA binding ^10^, ablated the ability of anillin to support junctional RhoA (Fig. 1g; Extended Data Fig. 3a,b) or stabilize either GFP-RhoA or GFP-RhoA^Q63L^ (Extended Data Fig. 3 f,g; Fig. 1 h,i). Since anillin has been implicated in cell division ^9^, we asked if RhoA stabilization also pertained during cytokinesis. Anillin KD destabilized GFP-RhoA^Q63L^ in both FRAP (Fig. 1m; Extended Data Fig.3i) and photoactivation (Fig. 1n; Extended Data Fig.3 j,k) experiments and stability was restored by anillin^WT^ but not by anillin^A740D/E758K^.

We then targeted the AH domain to specific cortical sites in anillin KD cells to test how RhoA scaffolding contributed to anillin-dependent contractilty. For the ZA, we fused AH to the N-terminus of α-catenin (AH-α-catenin) (Fig.2a; Extended Data Fig. 4a-c), using full-length α-catenin to avoid potential dominant-negative effects of α-catenin fragments; and for the cytokinetic furrow we used a previously-reported MRLC-AH fusion^10^. For comparison, we also targeted the high-affinity RhoA-binding domain of Rhotekin (rGBD). Binding of GTP-RhoA by the AH domain was sufficient to support RhoA activity to the ZA, being restored by AH-α-catenin but not by AH^A740D/E758K^-α-catenin (Fig. 2 b,c; Extended Data Fig. 4 c,d). Strikingly, AH-α-catenin also restored junctional levels of contractile effectors (Fig. 2 b,d; Extended Data Fig. 4c,e) and junctional tension itself (Fig.2e; Extended Data Fig. 4 f,g) to control levels. But this did not occur with AH^A740D/E758K^-α-catenin. Similarly, MRLC-AH, but not MRLC-AH^A740D/E758K^, largely restored the cytokinetic defects of anillin KD cells (Fig. 2f). In contrast, targeting the rGBD did not restore contractility to the ZA (Fig.2e; Extended Data Fig. 4 f), despite stabilizing RhoA as effectively as did AH-α-catenin (Extended Fig. 5 f-i). Nor was cytokinesis restored by an MRLC-rGBD fusion protein (Fig. 2f). The anillin AH domain alone was therefore sufficient to scaffold RhoA-dependent cell contractility at both the ZA and cytokinetic furrow, ruling out traditional multi-domain tether mechanisms.

Together, these findings present an apparent contradiction, since biochemical and structural studies have demonstrated that anillin binds to the same surface of GTP-RhoA as do effector proteins ^10,15,16^. Therefore, how could binding by anillin stabilise GTP-RhoA without sequestering it away from effectors and inhibiting signaling?

To explain this, we make contact with the notion of stochastic resetting ^17,18^, and propose that repeated, transient binding to anillin can increase the amount of time each GTP-RhoA molecule spends in its free (*i.e.*, non-anillin or -effector bound) state. This is only possible if, after unbinding from anillin, the inactivation rate of RhoA increases with time (a so-called non-Poisson process). Here, inactivation can then be thought of as being monitored by a stop-clock. The probability of inactivation (per unit time) increases with time unless there is an interruption, whereby the clock is reset. Transient binding of anillin constitutes precisely such an interruption, so that cycles of binding/unbinding repeatedly reset the inactivation “clock”, increasing the cortical residence time of free GTP-RhoA and hence the number of effector interactions. Importantly, this approach predicts that levels of RhoA-mediated contractile signalling can be tuned by modulating the local concentration of anillin, even if the level of RhoA is unchanged (Theoretical Supplement Fig. 1 a-g).

We developed a minimal cellular system to test our theory, by substituting AH for the cytoplasmic domain of the interleukin-2 receptor α-subunit (Tac, IL2R-AH; Fig. 3a). IL2R-AH was expressed in single MCF-7 cells along with a uniform amount of RhoA^Q63L^ mRNA, and the chimeric anillin construct concentrated into cortical patches by applying beads coated with α-Tac mAb (Fig. 3 a,b). As expected, IL2R-AH co-accumulated RhoA^Q63L^ in the cortical patches, but not if its ability to bind GTP-RhoA was ablated (IL2R-AH^A740D/E758K^, concentrated to a similar degree as IL2R-AH; Fig. 3 b,c; Extended data Fig. 5a). Furthermore, mDia1 co-accumulated in IL2R-AH, but not IL2R-AH^A740D/E758K^, patches (Fig. 3 d,e). Effector recruitment was confirmed using the ROCK1-GBD (Extended Data Fig. 5 b,c) and mDia1-GBD probes (Extended Data Fig. 5d), which presumably were responding to the presence of cortical GTP-RhoA, as they lack other protein-interaction domains. We then modulated the cortical concentration of IL2R-AH in the patches by coating the beads with different concentrations of α-Tac. FRAP experiments revealed that IL2R-AH stabilized GFP-RhoA^Q63L^ and the degree of stabilization increased with the density of IL2R-AH (Fig. 3f; Extended Fig. 5e). Strikingly, we found that the cortical recruitment of the effector mDia1 also increased with the amount of IL2R-AH (Fig. 3g), confirming the expectations of our resetting model.

**Figure 3.**
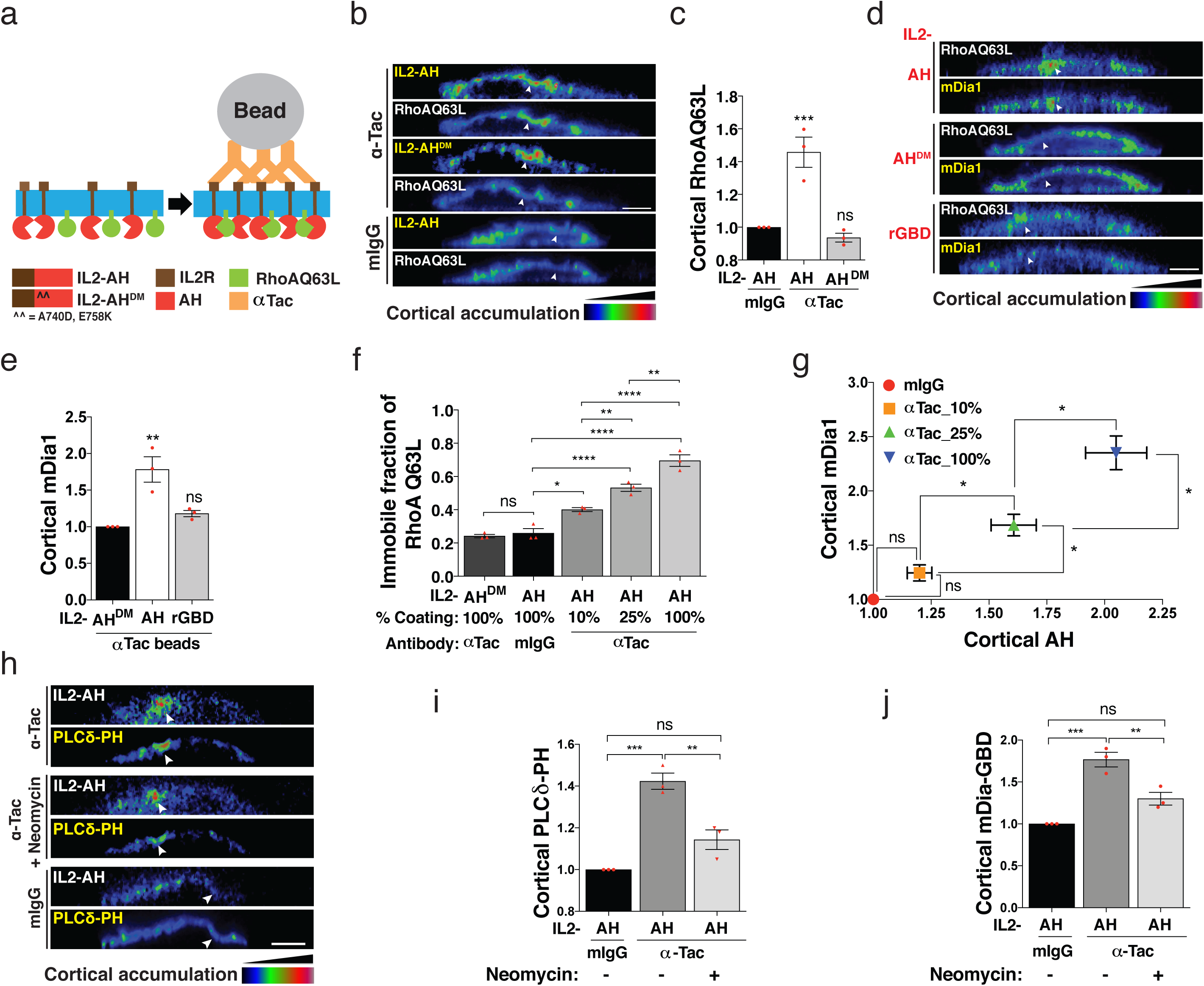
A non-tether kinetic model for scaffolding GTP-Rho by anillin. (**a**) Cartoon depicting the IL2-AH chimeric constructs and assay to co-cluster AH and RhoA on the cortex. (**b,c**) Clustering IL2-AH and RhoAQ63L with anti-tac beads. Representative heat map images (XZ views) of isolated MCF-7 cells co-expressing GFP-RhoAQ63L with either IL2-AH-Cherry or IL2-AH^A740D,E758K^-Cherry (AH^DM^) and overlaid with latex beads coated with anti-Tac or control mouse IgG. Arrows indicate the position of the latex bead on the cortex (**b**). Fluorescence intensity (FI) of cortical GFP-RhoA Q63L accumulated under beads (normalized to signal at the free cortex) (**c**). (**d,e**) Effect of clustering AH and rGBD domains on the recruitment of endogenous mDia1. Representative heat map images (XZ views) (**d**) and fluorescence intensity of mDia1 accumulated under beads (normalized to signal at the free cortex) (**e**). **(f,g**) Effect of clustering density of IL2R-AH on cortical stability of GFP-RhoA Q63L and recruitment of endogenous mDia1; immobile fractions from best-fit FRAP profiles (**f**); Fluorescence intensity of mDia1 and IL2-AH accumulated under beads (normalized to signal at the free cortex) with varied anti-tac coating (**g**). (**h-j**) Effect of clustering AH on cortical accumulation of membrane phospholipids (PIPs); Representative XZ images (**h**) and fluorescence intensity of PLCδPH (as a proxy for phosphotidylinositol 4,5 bisphosphate) (**i**) and GFP-mDia-GBD (**j**) underbeads coated with anti-tac and treated with neomycin. Data represent means ± s.e.m and n = 3 independent experiments. *P < 0.05, ** P < 0.01, ***P < 0.001, ****P < 0.0001; ns, not significant; One-way ANOVA with Dunnett’s multiple comparisons test (**c,e**) or Tukey’s multiple comparisons test (**f,g,i,j**); Scale bars, 5 μm.

Importantly, time-dependent rates (and their associated waiting-time distributions) typically arise from intermediate steps and cycles, as exemplified by the celebrated Hopfield-Ninio model of kinetic proofreading ^19–21^. This implied that anillin might affect some intermediate step in the process of RhoA inactivation. Given that anillin antagonized the dissociation of RhoA^Q63L^, we considered properties of the membrane that might influence this process. Notably, the AH domain has an atypical lipid-binding C2 motif ^10^ and IL2R-AH clusters accumulated phosphoinositide 4,5-P_2_ (PIP_2_) (Fig. 3h,i). As acidic phospholipids can antagonize RhoA dissociation both directly ^22^ and indirectly ^23^, we hypothesized that binding of anillin might facilitate the interaction of GTP-RhoA with a membrane environment that favoured its cortical retention. Indeed, blocking PIP_2_ access with neomycin significantly reduced the recruitment of RhoA effectors (tested using mDia1-GBD to eliminate potential contributions from the phospholipid-binding domain of mDia1) (Fig. 3j). Moreover, using a revised, fully Poissonian reaction scheme that explicitly incorporates the role of PIP_2_ as an inhibitor of membrane dissociation we predict that the level of effector recruitment is directly proportional to the level of anillin present at the membrane (Theoretical Supplement Fig. 1 h-k). This is consistent with our experimental findings (Fig. 3g), and supports the idea that antagonism of RhoA dissociation by concentrating membrane lipids may be instrumental for the AH domain to regulate GTP-RhoA signaling.

Finally, we sought the mechanism that controlled the cortical density of anillin, which our model predicted would determine how much it influenced RhoA signaling. Here, we noted that, although anillin is sensitive to RhoA (Fig. 4a,d), its junctional recruitment was most impaired when its NMII-binding domain was deleted (Extended Data Fig. 2 b,c). As NMII concentrates at sites where anillin is found, we postulated that NMII might constitute an additional cortical anchor for anillin. In support of this, the myosin inhibitor, blebbistatin, reduced junctional anillin (Extended Data Fig. 6a,b) and the stability of anillin in FRAP experiments was compromised when NMII-binding, but not F-actin binding, was disrupted (Extended Data Fig. 2 d,e). However, it was possible that these effects were indirect, because junctional RhoA signaling is compromised when NMII is blocked ^12,24^.

**Figure 4.**
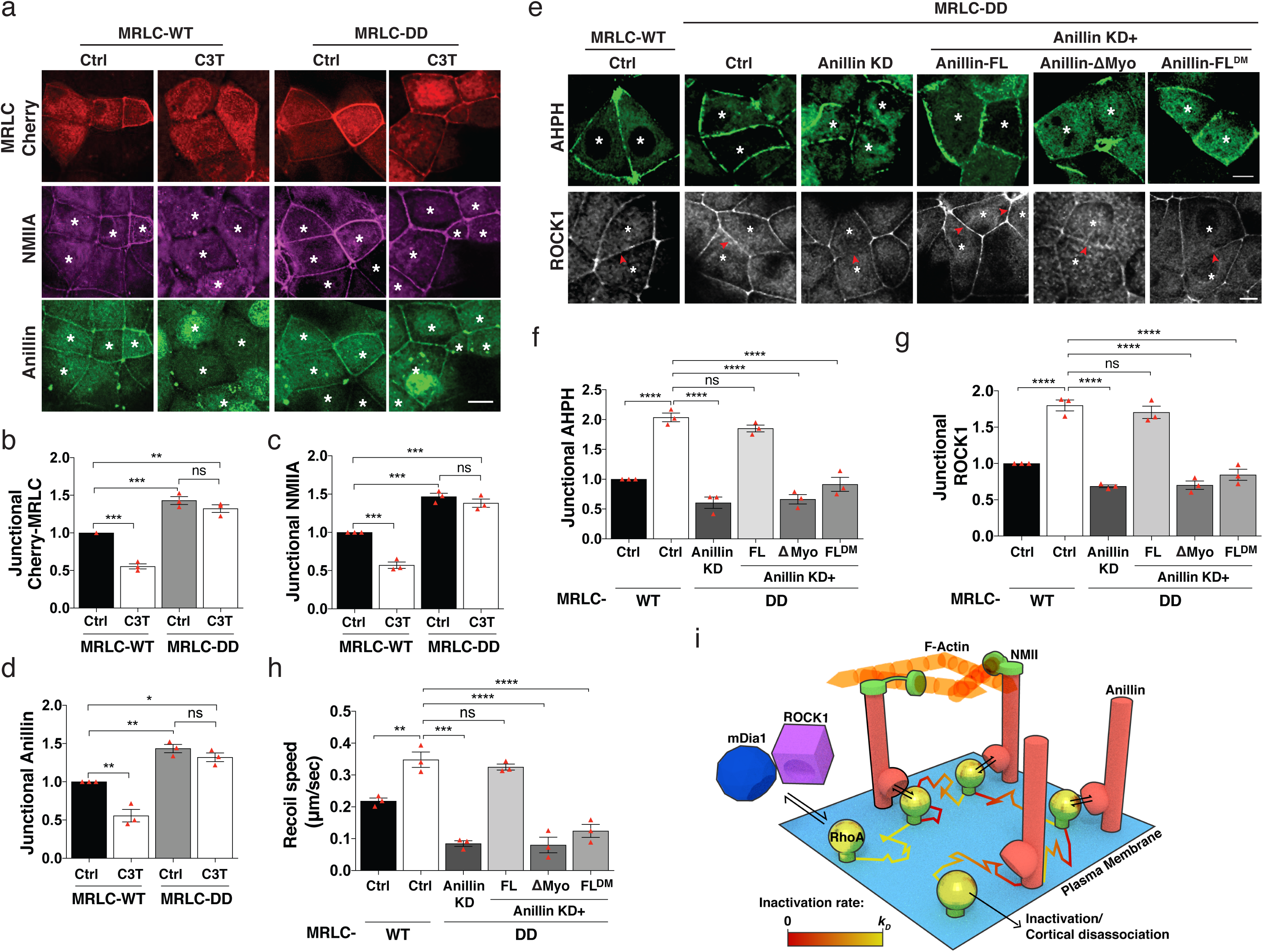
Cortical anchorage of anillin by Myosin II regulates RhoA signaling. (**a-d**) Myosin dependent localization of anillin; Representative images (**a**) and fluorescence intensity of cherry-MRLC (**b**), NMIIa (**c**) and anillin (**d**) at the ZA of cells expressing MRLC-WT or MRLC-DD and treated with C3 transferase (C3T). (**e-g**) Stabilized Myosin promotes RhoA signaling at ZA through anillin; Representative images (**e**) and quantified fluorescence intensity of AHPH (**f**) and ROCK1 (**g**) at the ZA of cells expressing MRLC-WT or co-expressing MRLC-DD with anillin shRNA (KD) or reconstituted anillin transgenes. Asterisks denote cells expressing the indicated transgene. Arrows indicate the homologous junctions that are quantified. (**h**) Junctional tension (initial recoil velocity) in cells expressing the indicated siRNA and transgenes. (**i**) Cartoon illustrating GTP-RhoA scaffolding by anillin. Free GTP-RhoA on membrane is able to interact with contractile effectors (mDia1 and ROCK1) so long as it does not undergo inactivation / cortical dissociation (whose rate is k_D_). Binding to Anillin blocks both inactivation and engagement with effectors. On un-binding, GTP-RhoA is again free to interact with effectors, however the rate of inactivation is now lowered, recovering to k_D_ with time. At sufficiently high density of anillin, repeated cycles of binding / unbinding can increase the residence time of free GTP-RhoA and hence interaction with effectors, until eventually the GTP-RhoA is inactivated. The cortical NMII network stabilizes and generates high density of anillin at the cortex which then provides a pathway for mechanochemical feedback from NMII to RhoA. Data represent means ± s.e.m and n = 3 independent experiments. *P < 0.05, ** P < 0.01, ***P < 0.001, ****P < 0.0001; ns, not significant; One-way ANOVA with Tukey’s multiple comparisons test; Scale bars, 10 μm.

Therefore, we developed a strategy to test if NMII can recruit anillin to junctions independently of RhoA. For this, we expressed a phosphomimetic transgene of MRLC (T18D/S19D; MRLC^DD^) that stabilized junctional GFP-NMIIA compared with a wild-type transgene (MRLC^WT^; Extended Data Fig. 6 c,d). Endogenous NMIIA persisted with MRLC^DD^ at junctions when RhoA was blocked with C3T (confirmed by loss of TCA-resistant RhoA staining; Fig. 4 a-c; Extended Data Fig. 6 f,g). In contrast, C3T displaced NMIIA from junctions in cells expressing MRLC^WT^ (Fig. 4 a-c). Thus, MRLC^DD^ could stabilize junctional NMII even when RhoA was inhibited. Importantly, MRLC^DD^ increased junctional anillin even in the presence of C3T (Fig. 4a,d; Extended Data Fig. 6 e), whereas junctional anillin was depleted in cells expressing MRLC^WT^ when RhoA was blocked. Thus, NMII could anchor and control the junctional concentration of anillin independently of RhoA.

Then we asked if stabilizing NMII modulated RhoA signaling via anillin. Indeed, MRLC^DD^ increased active RhoA at the ZA (Fig. 4 e,f; Extended Fig. 6 h,i) and increased the immobile fraction of GFP-RhoA^Q63L^ in FRAP experiments (Extended Data Fig. 6 j,k). This was accompanied by increased levels of the contractile effectors, ROCK1 and mDia1, and increased junctional tension (Fig. 4 e,g,h; Extended Data Fig.6 h, l-n). Anillin was necessary for these effects, as they did not occur in anillin KD cells, implying that MRLC^DD^ did not promote junctional contractility by a direct effect on NMII motor activity or via other pathways that can modulate RhoA ^12^. Furthermore, as predicted by our model, binding of RhoA by anillin was necessary for MRLC^DD^ to increase GTP-RhoA and contractility, as these effects did not occur in anillin KD cells reconstituted with anillin^A740D/E758K^. Nor were they restored if anillin was unable to bind NMII (anillin^ΔMyo^).

In conclusion, we propose that anillin functions as an NMII-anchored scaffold that promotes RhoA-dependent contractility by kinetic resetting (Fig 4i). In this model, cycles of unbinding and re-binding can increase the residence time of free, GTP-RhoA at the cortex, thus increasing its probability of engaging with effectors. For this, the process of RhoA inactivation must be non-Poissonian, implying that it includes some intermediate stage(s) that could be interrupted by anillin. Currently, our data with a GTP-locked RhoA suggest that one such intermediate factor is the content of acidic phospholipids in the local membrane environment that can antagonize RhoA dissociation ^22,23^. We postulate that anillin localizes PIP_2_, so that binding between anillin and GTP-RhoA increases the probability of interaction between PIP_2_ and GTP-RhoA. On unbinding from anillin, this lipid association facilitates membrane retention of GTP-RhoA, with the probability of cortical dissociation increasing with time (an inactivation “clock”). Cycles of binding / un-binding of anillin to GTP-RhoA may then repeatedly reset the clock, implying that the concentration of anillin can be used to tune the residence time of free GTP-RhoA, and hence interaction with effectors. Scaffolding by resetting can therefore be seen to be a regulatory analogue of kinetic proof-reading. In proofreading, a binding affinity, or time to unbinding, is encoded as a concentration. Here, the reverse is true: changing a concentration modifies a residence time (and hence effector engagement).

In this context, anchoring anillin by NMII might represent a post-activation mechanism to modulate contractile signaling that is orthogonal to the classical pathways that regulate RhoA signaling. Indeed, we found that stabilization of NMII enhanced junctional RhoA signaling via anillin, identifying both anillin and ROCK1 ^12^ as part of a feedback network that allows NMII to positively regulate RhoA signaling, its upstream regulator. Such positive feedback may account for the stable patterns of contractility found when epithelia assemble mature zonulae adherente ^11,12,24^, the exact adhesive zones where stable RhoA signaling coincides with the highest level of contractile tension.

More generally, scaffolding by repeated, transient binding may present an alternative paradigm for molecules that do not conform to the classical model of multi-domain tethering. Potentially this may apply to other GTPases as well as other signals whose functional outcomes depend on their dwell time in the active state.

## Acknowledgements

We thank our lab colleagues for much support and advice during the course of this work. This work was supported by grants (1037320, 1067405) and fellowships (1044041 [ASY] from the National Health and Medical Research Council of Australia, Australian Research Council (DP150101367, FT160100366 [GAG]), Queensland Cancer Council grants 1086857, 1128123), the Simons Foundation [RGM] and Human Frontiers Science Program grant RGP0023/2014). Optical microscopy was performed at the ACRF/IMB Cancer Biology Imaging Facility, established with the generous support of the Australian Cancer Research Foundation.

## Author contributions

SB and ASY conceived the project with conceptual and analytical input from GAG and NAH; SB designed and performed the experiments with technical assistance of MN and SV for cloning and western blotting respectively; S.B analysed the data. KBH and RGM developed the theory; and SB, KBH, RGM and ASY wrote the paper.

## EXTENDED DATA FIGURE LEGENDS

**Extended Data Figure 1:**
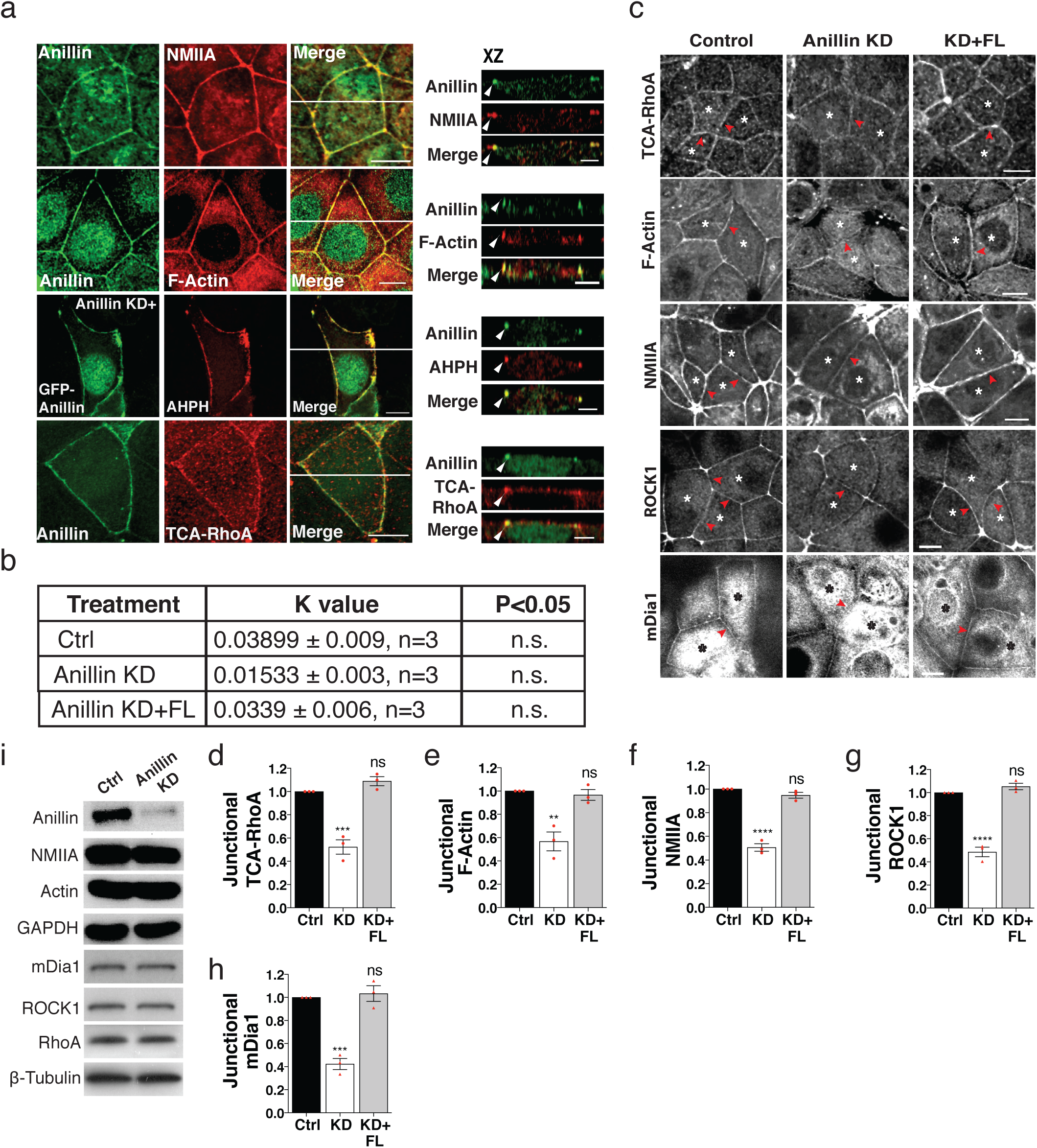
Related to Figure 1: Anillin binds and stabilizes active RhoA at sites of contractility. (**a**) Co-localization of endogenous anillin with NMIIA, F-actin, AHPH and TCA-RhoA. XZ views, taken at the indicated lines, show apical accumulation of anillin with NMIIA, F-actin, AHPH and TCA-RhoA. (**b**) Analysis of viscous drag from recoil measurements after laser ablation. The rate constant, k-value, for each junction was obtained after fitting the vertex displacement values to a mono exponential curve. To assess the influence of viscous drag on the initial recoil values used for tension measurements, the average k-values over several experiments were analysed for each condition. (**c-h**) Immunostaining of junctional proteins in MCF-7 cells expressing control shRNA, Anillin shRNA (KD), or Anillin shRNA reconstituted with full-length RNAi resistant Cherry-anillin). Representative images (**c**) and fluorescence intensity of junctional TCA-resistant RhoA (**d**), F-actin (**e**), NMIIA (**f**), ROCK1 (**g**) and mDia1 (**h**) at the ZA. Asterisks denote the cells expressing the indicated shRNA or transgene. Arrows indicate the homologous junctions that are quantified. (**i**) Immunoblot of anillin, NMIIA, actin, RhoA, ROCK1 and mDia1 in MCF-7 cells expressing control siRNA or anillin siRNA (KD). GAPDH and β-tubulin were loading controls. Data represent means ± s.e.m and n = 3 independent experiments. ** P < 0.01, ***P < 0.001; ****P < 0.0001; ns, not significant; One-way ANOVA with Dunnett’s multiple comparisons test. Scale bars, 10 μm; 5 μm for XZ views.

**Extended Data Figure 2:**
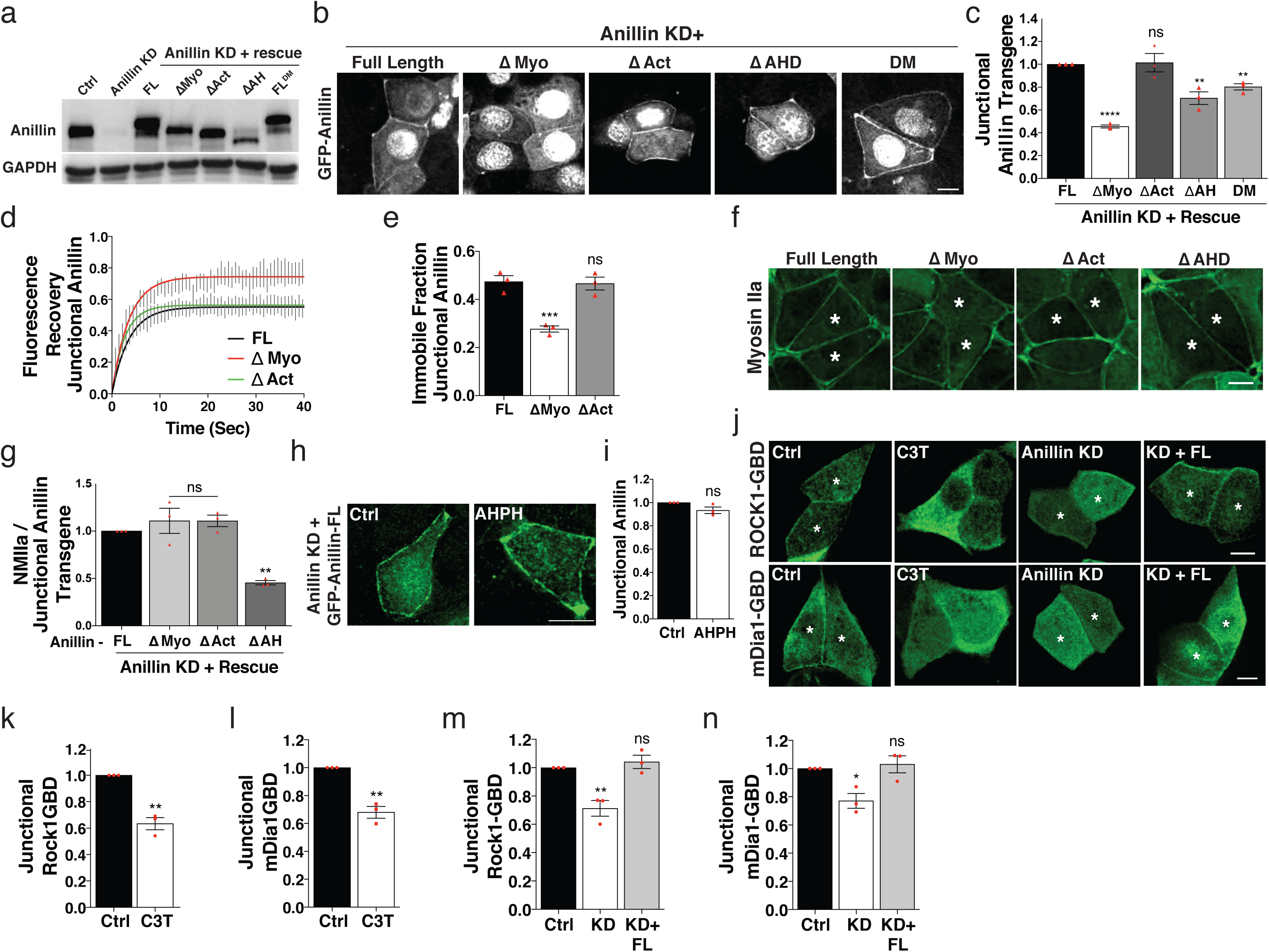
Related to Figure 1: Anillin binds and stabilizes active RhoA at sites of contractility. (**a**) Immunoblot of anillin in cells expressing anillin shRNA (KD) along with indicated RNAi resistant anillin transgenes. GAPDH served as loading control. (**b,c**) Representative images (**b**) and Junctional localization of full-length (FL) anillin and mutant transgenes normalized to expression levels (Junctional/cytoplasmic fluorescence intensity ratios) (**c**). (**d,e**) FRAP of anillin transgenes at the ZA; Best fit curves (**d**) and immobile fractions (**e**). (**f,g**) Junctional NMIIA levels in cells expressing anillin transgenes; Representative images (**f**) and fluorescence intensity of NMIIA normalized to junctional levels of anillin transgene (**g**). Asterisks denote cells expressing the indicated transgene. (**h,i**) Representative images (**h**) and fluorescence intensity of junctional anillin full-length transgene in cells expressing Cherry-AHPH (**i**). (**j-n**) Junctional localization of GFP tagged GTPase Binding Domain (GBD) of ROCK1 or mDia1. Representative images (**j**) and quantified junctional intensity of ROCK1-GBD and mDia1-GBD at ZA of cells treated with C3 transferase (1μg/ml) (**k,l**) or expressing anillin shRNA (KD) or reconstituted full-length anillin (**m,n**). Asterisks indicate cells expressing the transgene. Data represent means ± s.e.m and n = 3 independent experiments. *P < 0.05, ** P < 0.01, ***P < 0.001; ns, not significant; Student’s *t*-test (**i,k,l**), One-way ANOVA with Dunnett’s multiple comparisons test (**c,e,g,m,n**). Scale bars, 10 μm.

**Extended Data Figure 3:**
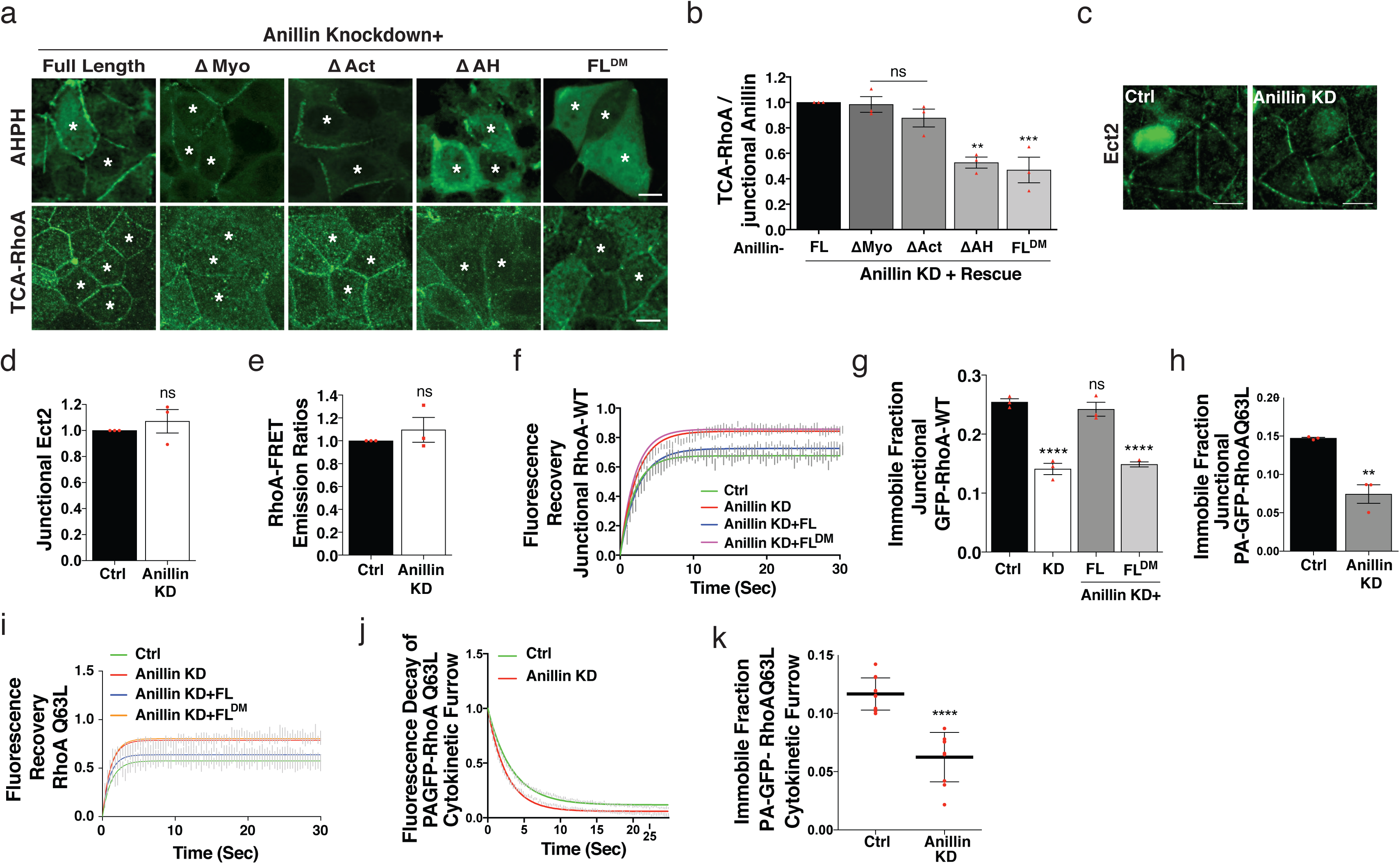
Related to Figure 1: Anillin binds and stabilizes active RhoA at sites of contractility. **(a-b)** Representative images of AHPH and TCA-RhoA (**a**) and fluorescence intensity of TCA-RhoA normalized to junctional levels of anillin transgene (**b**) at the ZA of cells reconstituted with anillin transgenes. (**c,d**) Immunostaining of Ect2 in cells depleted of Anillin (KD). Representative images (**c**) and fluorescence intensity at ZA (**d**). (**e**) RhoA FRET biosensor emission ratios measured at ZA in cells expressing control or Anillin siRNA. (**f,g**) FRAP of GFP-RhoA at the ZA in Anillin KD or reconstituted cells: best-fit curves (**f**) and immobile fractions (**g**). (**h**) Immobile fraction of photoactivated RhoAQ63L (PA-RhoAQ63L) at ZA in control or Anillin shRNA MCF-7 cells. (**i**) Best-fit curves of FRAP of GFP-RhoAQ63L at the cytokinetic furrow of Anillin KD or reconstituted cells. (**j,k**) Fluorescence decay (**j**) and immobile fraction (**k**) of photoactivated RhoAQ63L (PA-RhoAQ63L) at the cytokinetic furrow of control or Anillin shRNA MCF-7 cells. Data represent means ± s.e.m and n = 3 independent experiments except for (**k**) where data is means ± s.d. and n ≤11 cells. ** P < 0.01, ***P < 0.001, ****P < 0.0001; ns, not significant; Student’s *t*-test (**d,e,h,k**), One-way ANOVA with Dunnett’s multiple comparisons test (**b,g**). Scale bars, 10 μm.

**Extended Data Figure 4:**
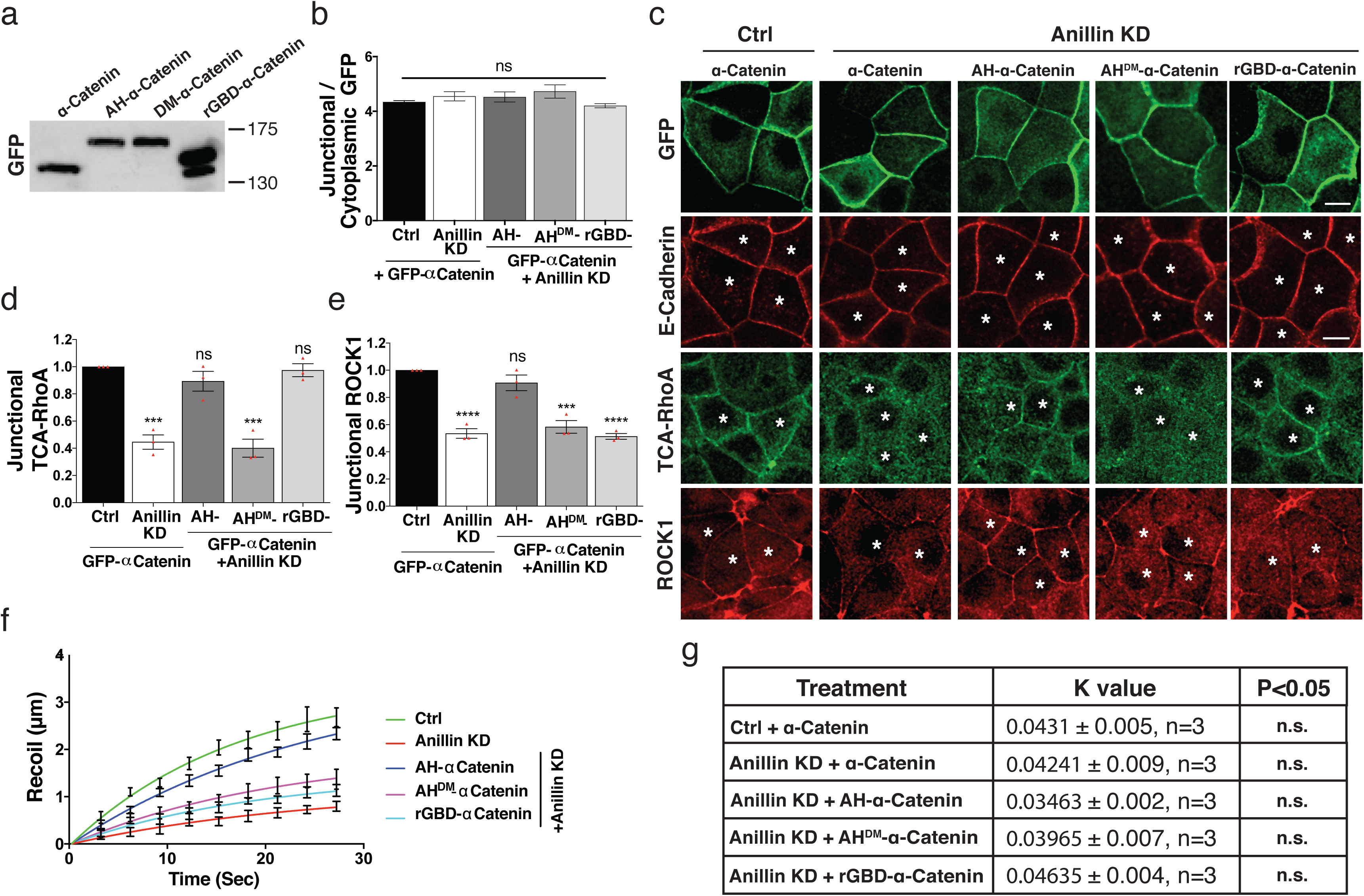
Related to Figure 2: The anillin AH domain is sufficient to support junctional contractility. (**a**) Immunoblot of GFP in cells expressing GFP tagged α-catenin chimeras. (**b-e**) GFP tagged α-catenin chimeras in MCF-7 cells. Fluorescence intensity of α-catenin chimeras normalized to cytoplasmic signal (**b**). Representative images of transgenes, E-cadherin, TCA-RhoA and ROCK1 (**c**) and fluorescence intensity of TCA-RhoA (**d**) and ROCK1 (**e**) at the ZA. Asterisks indicate cells expressing the chimeric transgenes. (**f,g**) Junctional tension in cells expressing the indicated siRNA and transgene. Recoil measurements of cell junctions after laser ablation (**f**). Analysis of viscous drag from recoil measurements after laser ablation (**g**). Data represent means ± s.e.m and n = 3 independent experiments. *P < 0.05, ***P < 0.001, ****P < 0.0001; ns, not significant; One-way ANOVA with Dunnett’s multiple comparisons test. Scale bars, 10 μm.

**Extended Data Figure 5:**
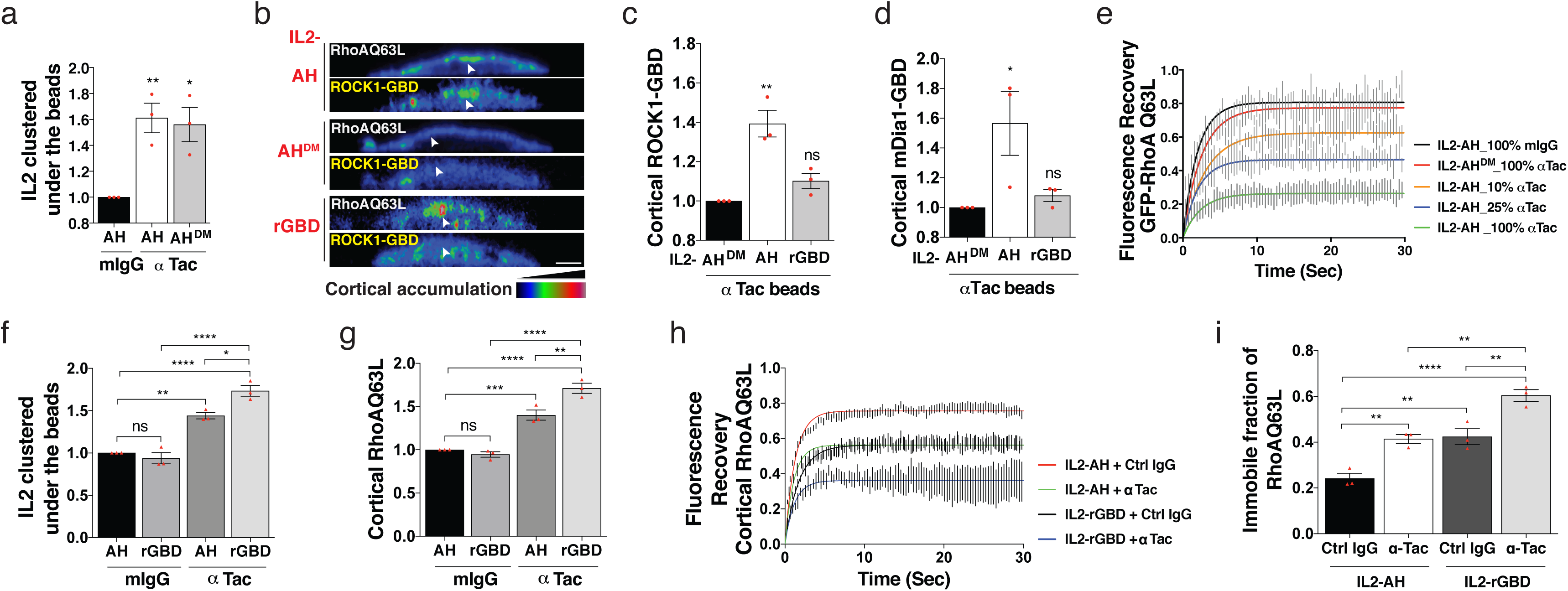
Related to Figure 3: A non-tether kinetic model for scaffolding GTP-Rho by anillin. (**a**) Clustering IL2 with anti-tac beads. Fluorescence intensity (FI) of cortical IL2 (-AH or - AH^DM^) accumulated under beads (normalized to signal at the free cortex). (**b-d**) Effect of clustering AH, AH^DM^ and rGBD domains on the recruitment of GFP-ROCK1-GBD and GFP-mDia1-GBD. Representative heat map images (XZ views) of ROCK1-GBD (**b**) and fluorescence intensity of ROCK1-GBD (**c**) and mDia1-GBD (**d**) accumulated under beads (normalized to signal at the free cortex). (**e**) Effect of clustering density of IL2R-AH on cortical stability of GFP-RhoA Q63L; best-fit FRAP profiles. (**f-i**) Clustering AH and rGBD domains with anti-Tac coated beads in isolated cells. Fluorescence intensity of IL2-AH-Cherry (**f**) and GFP-RhoA Q63L (**g**) accumulated under beads (normalized to signal at the free cortex). Effect of clustering AH and rGBD on cortical dynamics of GFP-RhoA Q63L, as measured by FRAP, in isolated cells: best-fit curves (**h**), immobile fractions (**i**). Data represent means ± s.e.m and n = 3 independent experiments. *P < 0.05, ** P < 0.01, ***P < 0.001, ****P < 0.0001; ns, not significant; One-way ANOVA with Dunnett’s multiple comparisons test (**a,c,d**) or Tukey’s multiple comparisons test (**f,g,i**). Scale bars, 5 μm.

**Extended Data Figure 6:**
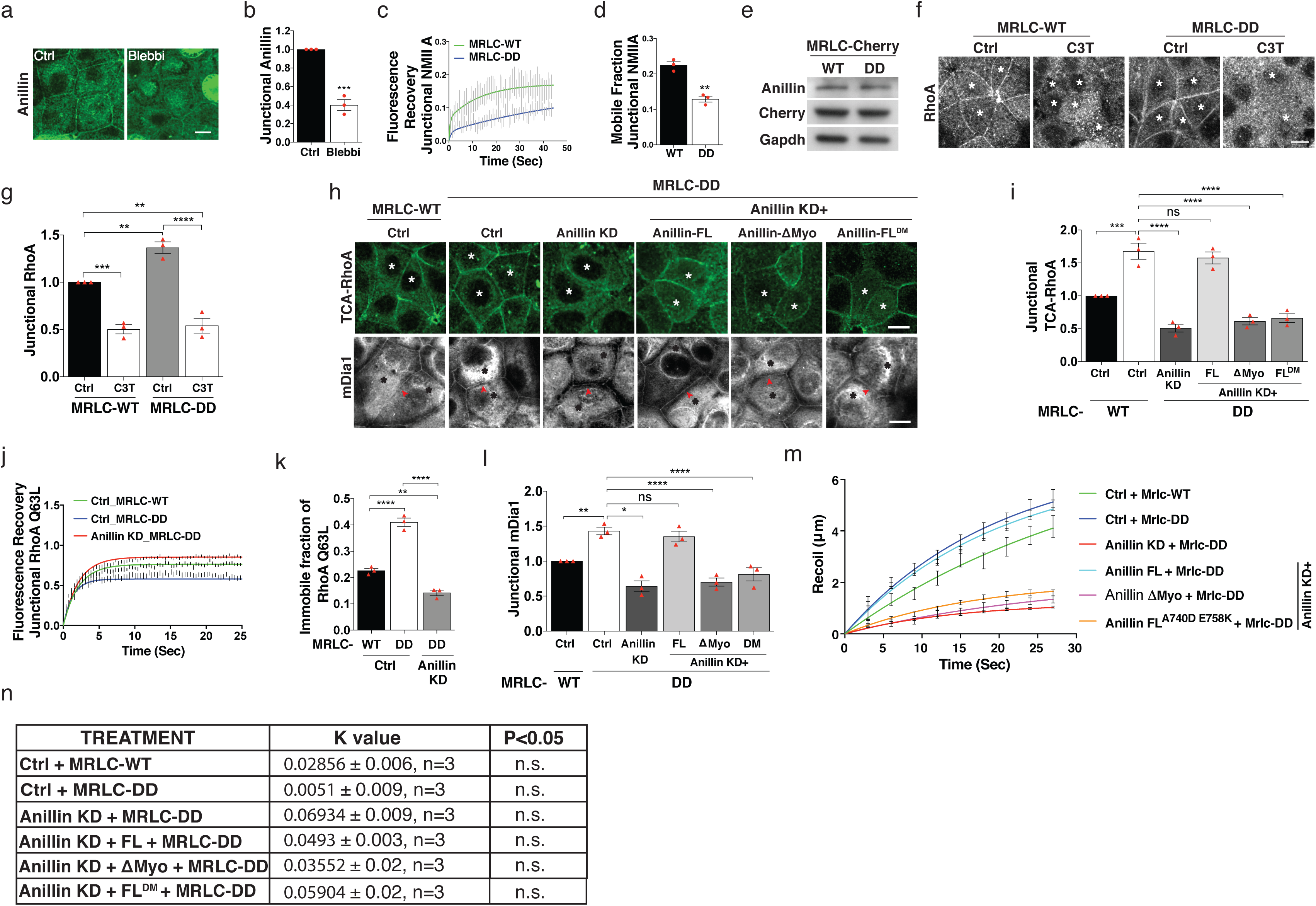
Related to Figure 4: Cortical anchorage of anillin by Myosin II regulates RhoA signaling. (**a,b**) Immunostaining of anillin in cells treated with blebbistatin. Representative images (**a**) and quantified fluorescence intensity at ZA (**b**). (**c,d**) FRAP of GFP-NMIIA at the ZA in cells expressing MRLC-WT or MRLC-DD: best-fit curves (**c**) and mobile fractions (**d**). (**e**) Immunoblot of anillin and mCherry in MCF-7 cells expressing mCherry tagged MRLC-WT or MRLC-DD. GAPDH serves as loading control. (**f,g**) Immunostaining of TCA-RhoA in cells expressing MRLC-WT or MRLC-DD and treated with C3 transferase (C3T). Representative confocal images (**f**) and quantified fluorescence intensity of junctional RhoA (**g**) (**h,i,l**) Representative images (**h**) and fluorescence intensity of TCA-RhoA (**i**) and mDia1 (**l**) at the ZA in cells expressing MRLC-WT or co-expressing MRLC-DD with anillin shRNA (KD) or reconstituted anillin transgenes. Asterisks denote cells expressing the indicated transgene. (**j,k**) FRAP of GFP-RhoA Q63L at the ZA in cells expressing MRLC-WT, MRLC-DD or co-expressing MRLC-DD with Anillin shRNA (KD): best-fit curves (**j**) and immobile fractions (**k**). (**m,n**) Recoil measurements of cell junctions after laser ablation of junctions of cells expressing MRLC-WT or co-expressing MRLC-DD with Anillin shRNA (KD) or reconstituted Anillin transgenes (**m**). Analysis of viscous drag from recoil measurements after laser ablation (**n**). Data represent means ± s.e.m and n = 3 independent experiments. *P < 0.05, ** P < 0.01, ***P < 0.001, ****P < 0.0001; ns, not significant; Student’s *t*-test (**b,d**) One-way ANOVA with Tukey’s multiple comparisons test (**g,i,k,l**). Scale bars, 10 μm.

## EXTENDED DATA MOVIE CAPTIONS

**Movie 1:** 3D reconstruction of MCF-7 immunostained for Anillin. Anillin exhibits predominant localization to apical junctions and nuclei.

**Movie 2:** Photoactivation of junctional RhoAQ63L. Fluorescence decay of photoactivated RhoAQ63L at apical junctions shows rapid loss of fluorescence in control and Anillin KD cells with no visible lateral dispersion.

