## Supplementary Materials for "Scaffolding of RhoA contractile signaling by anillin: a regulatory analogue of kinetic proofreading"

### SUPPLEMENTAL MATERIAL

#### A. Supplemental methods and materials.

**Cell Culture and Drug treatment:** MCF-7 cells were obtained from ATCC and were cultured in DMEM supplemented with 10% FBS, 1% L-Glutamine, 1% pencillin/streptomycin. The cell line was routinely tested for mycoplasma and maintained in a low dose of plasmocin (Invitrogen). Forty-eight hours post seeding, cells were transfected with plasmid DNA or SiRNA, at 60-70% confluency, with Lipofectamine 3000 or RNAimax (Invitrogen) respectively, as per manufacturers' instructions and analyzed after 24 hours. For live cell imaging experiments, cells were cultured on No. 1.5, 29 mm glass bottom dishes (Shengyou Biotechnology) and maintained in Hank's balanced salt solution supplemented with 5% FBS, 15 mM HEPES at pH 7.4, 10 mM D-Glucose and 2 mM Cacl2. To inhibit RhoA activity, cells were treated with 1 µg/ml of cell-permeable RhoA inhibitor (C3-T based; no. CT04-A, Jomar Bioscience) for 2–3 hours. To inhibit myosin, cells were treated with blebbistatin (catalogue no. US1203390-5MG) at a final concentration of 100 µM for 3 hours. To block phospholipids, Neomycin (catalogue no. N6386-25G) was used at 2 mM for 15 minutes.

**Plasmids, SiRNA and ShRNA:** GFP-Anillin and GFP-AHPH were a gift from M.Glotzer (University of Chicago) and was described previously[1]. mCherry-tagged anillin was generated by replacing GFP with mCherry using AgeI and BsrGI restriction enzyme sites. The shRNA targeting anillin [1] and GFP-Anillin were cloned into the lentivirus vector lentilox pLL5.0, a gift from J.Bear (UNC Chapel Hill, USA). The GFP- and Cherry-tagged Anillin$^{A740D,E758K}$ constructs were generated using the Quick Change II Site Directed Mutagenesis Kit as per the manufacturer's protocol. Deletion mutants of anillin (Δ Myosin Binding, Δ Actin binding, Δ Anillin Homology domain) were generated by assembling the PCR amplified fragments into SacII/SbfI restriction digested pLL5.0 containing anillin ShRNA using the Infusion cloning Kit (Clontech); primers for these are listed in the Supplemental table., mCherry-AHPH was generated by replacing GFP with mCherry using AgeI and BsrgI restriction enzyme sites. GFP-AHPH$^{A740D,E758K}$ is described previously [2]. GFP-tagged mouse mDia1 was a kind gift from A. Bershadsky (Mechanobiology Institute). FLAG-tag mouse ROCK1 was a kind gift from M. Samuel, (Centre for Cancer Biology, University of South Australia, Australia). GFP-Rock1-GBD (840-110 aa) and GFP-mDia1-GBD (1-305 aa) were generated by PCR amplifying these fragments from Flag-ROCK1 and GFP-mDia1 plasmids and cloned into pEGFP-C1 (Clontech) vector using Xho1/BamH1 and Xho1/EcoR1 sites respectively. pCAGGS-Raichu-RhoA-CR was obtained from Addgene (No. 40258). GFP-RhoA-WT and GFP-RhoAQ63L were purchased from Addgene (No. 12965 and No. 12968 respectively). mCherry-RhoAQ63L was generated by replacing GFP in GFP-RhoAQ63L with PCR amplified mCherry using the restriction enzymes HindIII/EcoR1. Photoactivable RhoAQ63L was generated by subcloning mRFP-PA-GFP from mRFP–PA-GFP–actin plasmid (a gift from G. Charras, UCL, UK) into GFP-RhoAQ63L plasmid digested with HindIII and Bsrg1. Both, GFP-RhoAQ63L and mRFP-PAGFP were blunted by Quick blunting kit (NEB) after restriction digestion with HindIII and AgeI respectively. pCMV-IL2R-hECD tail is a gift from Cara Gottardi (Feinberg School of Medicine, Northwestern University). IL2-Cherry was generated by cloning the PCR amplified extracellular and transmembrane domains of IL2 from the plasmid, pCMV-IL2R-hECD tail, into the pmCherry-N1 (Clontech) using the NheI/AgeI restriction sites. IL2-AH, IL2-AH$^{A740D,E758K}$ and IL2-rGBD were generated by assembling the PCR fragments of IL2 with AH or AH$^{A740D,E758K}$ or rGBD into pmCherry-N1 vector linearized with NheI and AgeI. pCS2-EGFP-rGBD was a kind gift from William Bement (University of Wisconsin-Madison). IL2-AH-2A-CherryRhoAQ63L, IL2-AH$^{A740D,E758K}$-2A-CherryRhoAQ63L and IL2-rGBD-2A-CherryRhoAQ63L were generated by assembling the PCR amplified 2A-Cherry-RhoAQ63L into the Cherry-excised IL2-AH-Cherry, IL2-AH$^{A740D,E758K}$-Cherry and IL2-rGBD-Cherry constructs using the AgeI/NotI restriction sites and Infusion Cloning kit. AH-GFP-αCatenin, AH$^{A740D,E758K}$-GFP-αCatenin and rGBD-GFP- αCatenin were generated by cloning synthesized AH, AH$^{A740D,E758K}$ and rGBD fragments into GFP-αCatenin plasmid using the NheI/AgeI restriction sites. These DNA fragments were synthesized from GenScript, USA. siRNA against the coding region of human anillin (UUUAUUCAAAGAGGCAUCGCCAUCC; Invitrogen, USA) were described earlier [1]. MRLC1-GFP and MRLC1-T18D, S19D-GFP were obtained from Addgene (No. 35682 and No. 35680). MRLC1-Cherry and MRLC1-T18D, S19D-GFP were generated by replacing GFP with Cherry using AgeI/BsrGI restriction enzymes. GFP-PLCδPH is a gift from Mark A. Lemmon (University of Pennsylvania and Yale university). The primer sequences used to clone these constructs are listed in Supplemental Table 1.

**Antibodies:** Anillin antibody raised in rabbits was a kind gift from Micheal Glotzer (University of Chicago) and was used at 1:200 for immunofluorescence and 1:1000 for immuoblotting. Other primary antibodies that were used were: mouse monoclonal antibody (mAb) HECD-1 against the ectodomain of E-cadherin (1:50; a gift from P. Wheelock, University of Nebraska, Omaha, Nebraska, USA; with the permission of M. Takeichi); rabbit polyclonal antibody (pAb) for non-muscle myosin IIA heavy chain (1:1,000; no. PRB-440P, Covance); rabbit pAb for non-muscle myosin IIB heavy chain (1:1,000; no. PRB-445P, Covance); rabbit pAb (1:1,000; no. A-6455) and mouse mAb (1:100; no. A-11120, clone 3E6) against GFP (Molecular Probes/Invitrogen); mouse mAbs against RhoA (1:100; Santa Cruz Biotechnology, clone 26C4, no. sc418); rabbit pAb against Ect2 (1:50; No. 07-1364; Millipore); rabbit pAb against ROCK1 (1:300, no. AB134181, Abcam); mouse mAb against GFP (no. 11814460001, Roche); mouse mAb against actin (1: 200, No. MAB1501, Millipore); rat mAb E-cadherin (1:1000, No. 13-1900, Invitrogen); and mouse mAb against IL2 (ab86897, Clone 7G7B6, Abcam). Species-specific secondary antibodies were used that are conjugated with AlexaFluor 488, 546 or 647 (1:500, Invitrogen) for immunofluorescence or with horseradish peroxidase (1:5000, Bio-Rad Laboratories) for immunoblotting.

**Immunoblotting**: Cells were lysed and protein samples were collected in lysis buffer (50 mM Tris-HCl, pH=7.4, 150 mM NaCl, 1% NP40, proteinase inhibitor cocktail [Roche, Cat#138467]). Samples were resolved on SDS-PAGE and transferred onto nitrocellulose membranes. After blocking in 5% non-fat milk in 0.1% Tween TBS, the membranes were incubated with the primary antibody for 1 hour at room temperature followed by washing with TBS containing 0.1% Tween 20. Following incubation with species-specific HRP conjugated secondary antibodies (1 hour at room temperature) signals were detected by Enhanced chemiluminescence (Pierce).

**Immunofluorescence microscopy and image analysis:** Cells were fixed with ice-cold methanol for 5 min at -20$^0$C or with 4% paraformaldehyde in cytoskeleton stabilization buffer (10 mM PIPES at pH 6.8, 100 mM KCl, 300 mM sucrose, 2 mM EGTA and 2 mM

MgCl2) on ice for 15 min or with freshly prepared 10% TCA in distilled water on ice for 15 min (for RhoA staining). TCA-fixed cells were subsequently washed three times with 30 mM glycine. Both PFA and TCA fixed cells were permeabilized with 0.25% Triton X-100 for 5 min at room temperature, followed by blocking with 5% fat free milk and incubation with primary antibodies (1h at room temperature or over night at $4^0$ C). The coverslips were washed with PBS and incubated with secondary antibodies for 1 hour at room temperature and coverslips were mounted on glass slides with Prolong Gold mountant (ThermoFisher). Confocal images were acquired on a Zeiss 510 or a Zeiss 710 Meta laser-scanning confocal microscope with a 63X PlanApo 1.4 NA objective.

Quantification of junctional fluorescence was performed by the line scan function of ImageJ software (NIH). A line of 30 pixel width and 10 µm length was drawn orthogonal to randomly chosen homotypic apical junctions (i.e. junctions between cells that are manipulated). For each independent experiment, 25 junctions were quantified for each individual conditions. The intensity value along the line was recorded with the Plot Profile feature of ImageJ and the peak intensity value round the center of the line was averaged across junctions. To quantitate junctional accumulation of various transgenes and AHPH, the peak intensity values were normalized to the average intensity of pixels in the cytoplasm. Contrast adjustment and *Z*-projections of raw data images were performed with ImageJ. Representative images were prepared by maximum projection of apical confocal stacks. For representation, raw images were processed in ImageJ by applying a median filter of one pixel radius. Where necessary, background subtraction was performed using the rolling-ball background subtraction function in ImageJ. Control and test images were processed identically.

**Quantification of bi nucleated cells:** Cells expressing the specific GFP tagged transgene (MRLC-GFP or MRLC-GFP fused with AH, $AH^{DM}$ or rGBD) were fixed with PFA and immunostained for GFP and DAPI. The number of bi/multi nucleated cells were recorded for 50 cells expressing the transgene for each condition per experiment using a 63X PlanApo 1.4 NA objective on a bright field microscope.

**Fluorescence Recovery After Photobleaching (FRAP) and PhotoActivation:** FRAP was performed on a LSM 510 Meta Zeiss confocal microscope with a Ti:Sapphire multiphoton laser (Chameleon Ultra, Coherent Scientific) or a Zeiss LSM 710 Meta confocal microscope with a Mai Tai (Spectra physics) multiphoton laser equipped with a $37^0$ C heating stage. MCF-7 cells were transfected with GFP-Anillin plasmids (WT or indicated mutants) or GFP-RhoA (WT or Q63L) along with desired expression plasmid. Images (57.14 × 57.14 microns) were acquired with a 63X,1.4NA Plan Apo objective and 3× zoom, A constant ROI was drawn in the centre of the cell–cell contact and was bleached to 70–80% with a 790 nm Chameleon laser at 38% power (one iteration) or 810nm MaiTai laser at 20% power (two iterations). Time series images were acquired before (10 frames, ~ 380 ms) and after bleaching (909 frames, 35.15 s). The mean fluorescence intensity values in the bleached area F(t) were analysed with ImageJ using a constant ROI. The residual intensity in the ROI of the first frame after bleaching F(0), was substracted from the mean intensity values F(t) and the average pre-bleach intensities F(i) in the ROI. Fluorescence intensities were normalized to the average pre-bleach values and fitted to a mono-exponential function in the GraphPad Prism software to calculate the Mobile fraction and half-time of recovery.

For photoactivation, MCF-7 cells were transfected with RFP-PAGFP-RhoAQ63L along with RhoA siRNA against the UTR region and siRNA against anillin. Images (53.98

× 53.98 microns) were acquired on a LSM 710 Meta Zeiss confocal microscope with a $37^0$ C heated chamber, 63× objective (1.4NA oil Plan Apochromat immersion lens) and 3× zoom. Photoactivation of PAGFP-RhoAQ63L with in a constant ROI at the apical cell-cell contact was achieved by Mai Tai multiphoton laser (2000 mW) at 810 nm 18 % power (one Iteration). Time series images were acquired before (5 frames, ~ 157 ms) and after photoactivation (1195 frames, 37.53 s). The mean fluorescence intensity values in a constant ROI within the photoactivated region were obtained by ImageJ for all time points. The fluorescence intensities were normalized to the first time point after photoactivation after correcting for the average intensity value before photoactivation and were fitted to a one-phase decay function in GraphPad prism software. For each independent experiment, FRAP from 8 junctions (in which both cells express the shRNA or reconstituted transgene) was measured for each individual condition.

**Junctional recoil measurements by laser ablation.** MCF-7 cells were transduced with lentivirus encoding E-cadherin– EGFP and transfected with desired plasmids or the various GFP-α-catenin constructs. Cells were processed for experiments 24h post transfection. Recoil measurements were performed on Zeiss LSM 510 Meta or LSM 710 Meta confocal microscopes equipped with $37^0$ C heating stages. Images (38.56 x 38.56 µm) were acquired with a 63x objective (1.4 NA oil Plan Apochromat immersion lens) with 3x zoom. A constant ROI, at the center of the cell–cell contact was ablated with using two-photon lasers (Chameleon Ultra, Coherent Scientific or MaiTai, Spectra-Physics) tuned at 790 nm (30 iterations,) and 25% transmission for Chameleon laser or at 810 nm (10 iterations), 15% transmission for Mai Tai laser. Time-lapse images were acquired before (2 frames) and after (7 frames) ablation at an interval of 3 or 6 seconds. The retraction of junctions over time was tracked by tracking function in ImageJ. The mean values of displacement of vertices were plotted over time and fitted to a mono exponential function in the GraphPad Prism software to calculate the initial recoil speed and rate constant as described previously [2-4]. For each independent experiment, recoil from 8 junctions (in which both cells express the shRNA or reconstituted transgene) was measured for each individual condition.

**FRET microscopy.** FRET measurements and analysis were performed 24 h after transfection as described previously [2,3]. Briefly, live-cell imaging was performed at $37^0$ C on an LSM710 Zeiss confocal microscope equipped with a 63x oil immersion objective (Plan Apochromat 1.4 NA, Zeiss) and GaAsP (BIG) detectors. Images were acquired by sequential line acquisition. The acceptor (A) channel was imaged using a 514 nm laser line for excitation and emission was collected in the acceptor emission range (BP 530–590 nm). Donor and FRET channels were acquired using a 458 nm laser line and emission was collected in the donor emission region (BP 470–490 nm) and acceptor emission region (BP 530–590 nm), respectively. The images were processed by ImageJ software. The acceptor images were used to create a mask of the junctional region to analyze FRET efficiency in this region using ImageJ. The FRET ratio (FRET/acceptor) was calculated as described previously by a custom MatLab routine [2,3]. For each independent experiment, FRET measurements were performed on 25 junctions for each individual condition.

**Clustering IL2 chimeras on cell membranes:** MCF-7 cells were co-transfected with the desired IL2 chimera plasmid along with any of these plasmids; GFP-RhoA-Q63L, GFP-ROCK1-GBD, GFP-mDia1-GBD. Post transfection (24 h), cells were overlaid with anti IL2 coated beads (latex-Spherotech #MPFc-30-5 or dynabeads-Thermofischer #11201D) for 15- 20 minutes before fixing or live imaging. To identify the latex beads on the cell surface, Z stacks of bright-field images were acquired along with fluorescence images with a Zeiss 710 Meta laser-scanning confocal microscope with a 63x PlanApo 1.4 NA objective. For FRAP, a constant ROI was marked and bleached on the cell membrane just below the anti IL2 coated dynabeads. Spatially define bleaching in the Z axis was achieved by a Chameleon multiphoton laser tuned at 710 nm or Mai Tai laser tuned at 810 nm. Generally 1µg of Anti IL2 or mIGg was allowed to bind to 50 µl of dynabeads or 150 µl of Latex beads in 0.5% BSA c for 30 minutes at room temperature and stored at $4^0$ C until further use. The beads were coated with antibodies 2 hours prior to experiment. To saturate the Anti IL2 coating on beads, 50 µl dynabeads or latex beads were allowed to bind to 4 µg of Anti IL2 or mIGg (100%). The Anti-IL2 coating on beads was varied (10% and 25%) by changing the amount of Anti-IL2 (1µg and 0.4 µg respectively) and using mIgG to maintain a constant amount of IgG that was incubated with beads. For quantifying cortical accumulation of IL2 probes, RhoAQ63L and Effectors (mDia, ROCK1-GBD and mDia-GBD), the cells with overlaid beads were immunostained for respective probes and proteins and Z stack images (Fluorescence and DIC) were acquired on confocal microscope (as described in immunofluorescence and image analysis section). The XZ profiles of the Z stack images were generated using ImageJ software and raw fluorescence intensity under the beads was measured using a constant ROI of 4 pixel width for 25 beads for each condition for independent experiment.

**Statistics and repeatability of the experiments**: Detailed statistical analysis for individual experiments are listed in Supplementary Table 2. This table contains the detailed statistical parameters generated for each experiment (also mentioned in the corresponding figure legend); this includes the statistical test performed and number of independent experiments. Throughout the text and as mentioned in the corresponding figure legends, n represents the number of independent experiments except for FRAP and photoactivation experiments at cytokinetic furrow where n represents number of cells/observations. To be precise, n defines the number of times the experiment was repeated independently at different times thus ensuring that it accounts for the variability of the biological process. For each assay, the number of observations within an independent experiment is mentioned in methods. To compare two or more groups, we used Student's t-test or one-way ANOVA, respectively. Moreover, when one-way ANOVA was used, we applied corrections for multiple comparisons depending on whether data were compared with the control group alone (Dunnett's) or multiple comparisons (Tukey's) between all the groups were made. This correction is mentioned in the corresponding figure legends. GraphPad Prism 6 was used to determine the P values and perform all statistical analysis. To determine the sample size for our assays, standard deviation was calculated from initial trials and used to determine the sample size based on confidence interval calculations at confidence levels of 95%. The experiments were not randomized, and the investigators were not blinded to allocation during experiments and outcome assessment. However, a third party blinded to our hypothesis analyzed the junctional recoil and immunofluorescence of effector recruitment (Fig. 2e; Fig 4h and Extended Data Figure 5 b-d). All representative images and videos were observed in three independent experiments (see Supplementary Table 2). For Western blots confirming knockdown and reconstitution of various transgenes (Extended data Fig. 2a) were performed twice for this study. The control and treated group of cells came from the same source and were processed at the same time, in a similar fashion and also imaged on the same day. The concentration and duration of drug treatments were rigorously established and kept constant during different experiments. For experiments related to AHPH and other GBD probes (ROCK1 and mDia1) transfection conditions were standardized to achieve the optimal level of expression.

Code availability. The MATLAB scripts for FRET analysis as well as the codes for computational simulations are available on request.

**SUPPLEMENTAL TABLE 1:**

| No. | Construct/Primer Name | Sequence (5'-3') |
|---|---|---|
| | **ANLN MUTANTS** | |
| 1 | PLL-SacII-GFP-F | AGTCGACGGTACCGCGGATGGTGAGCAAGGGCGAGGAGC |
| 2 | PLL-SbfI-Anln-R | CGACGAATTGCCTGCAGGTTAAGGCTTTCCAATAGGTTTGTAGC |
| 3 | Del Myo-Fragment 1-R | CTGCTTAACACTGCTTGCAAGTTTTTGCATACGTGTTTTAACTGAGG |
| 4 | Del Myo-Fragment 2-F | AGCAGTGTTAAGCAGGAAGCTACATTCTGTTCCC |
| 5 | Del Act-Fragment 1-R | TTGTGCTAAATGGGTGCTATTGATCCTAGCAGATGCTCCACTTGC |
| 6 | Del Act-Fragment 2-F | ACCCATTTAGCACAACAGCTCAAGCAGGAACG |
| 7 | Del AHD-Fragment 1-R | TTCAACACTGGAATTTGGTTTTGGACTTTCATCACTTCTGCTGG |
| 8 | Del AHD-Fragment 2-F | AATTCCAGTGTTGAAGAAAGAGGTTTTCTAACC |
| 9 | Del PH-Fragment 1-R | CGACGAATTGCCTGCAGGTTACACTTGACATTTTATTTTTAAATAAATATGACCTTCC |
| | **IL2-CHERRY** | |
| 10 | IL2Ra-NheI-F | GCTAGCATGGATTCATACCTGCTGATGTGGGG |
| 11 | IL2Ra-AgeI-R | AAACCGGTATCTTCCAGGTGAGCCCACTCAGG |
| | **IL2-AH AND IL2-DM** | |
| 12 | IL2Ra-vector Overlap-F | CGTCAGATCCGCTAGATGGATTCATACCTGCTGATGTGGGG |
| 13 | IL2Ra-AH Overlap-R | TCTTTGGAATTTTCCCTTCCAGGTGAGCCCACTCAGG |
| 14 | AH-F | GGGCTCACCTGGAAGGGAAAATTCCAAAGAACTCGTGTCCCTCGAGC |
| 15 | AH-Cherry overlap-R | CCATGGTGGCGACCGGTCGCACTTGACATTTTATTTTTAAATAAATATGACCTTCCAAAGAAGATAAAAAGGGG |
| | **IL2-rGBD-Cherry** | |
| 16 | XhoI-IL2-Fwd | TTCTCGAGATGGATTCATACCTGCTGATGTGGGG |
| 17 | AgeI-rGBD-Rvs | AAACCGGTGCGCCTGTCTTCTCCAGCACCTG |
| | **Cherry-RhoAQ63L** | |
| 18 | HindIII Cherry Fwd | TTAAGCTTGCCACCATGGTGAGCAAGGGCGAGGAG |
| 19 | EcoR1 Cherry Rvs | AAGAATTCCTTGTACAGCTCGTCCATGCC |
| | **IL2-AH and DM-2A-CherryRhoAQ63L** | |
| 20 | 2Acherry-F1 | GAAAACCCTGGCCCCGCCCCAGGATCCAAGCTTATGGTGAGCAAGGGCGAGGAGG |
| 21 | 2Acherry-F2 | AGGGGAAGCCTGCTCACCTGCGGCGACGTGGAGGAAAACCCTGGCCCCGCCCC |
| 22 | 2Acherry-F3 | ATAAAATGTCAAGTGCGACCGGTCGAGGGCAGGGGAAGCCTGCTCACCTGCGG |
| 23 | 2Acherry RhoA-R | TGATCTAGAGTCGCGGCCGCTCACAAGACAAGGCAACCAGATTTTTTCTTCCCACGTC |
| | **IL2-rGBD-2A-CherryRhoAQ63L** | |
| 24 | GFP n Cherry Infusion 2A-F | CCCCGCCCCAGGATCCAAGCTTATGGTGAGCAAGGGCGAGGAG |
| 25 | RhoA Q63L Infusion R | GTTATCTAGATCCGGTGGATCCTCACAAGACAAGGCAACCAGATTTTTTCTTCC |
| 26 | IL2-rGBD-2A | GCTCACCTGGAAGATACCGGTGAGTCGGATCACAGTGGC |

| | | |
|---|---|---|
| | (Synthesized) | CAGGGGCTCCGCCCTGGAGATGGAGTTCAAACGCGGCCGCTTCCGACTTAGTTTCTTCAGCGAGTCGCCGGAGGACACAGAGCTGCAGAGGAAACTAGATCATGAGATCCGGATGAGGGATGGGGCCTGCAAGCTGCTGGCAGCCTGCTCCCAGCGAGAGCAGGCTCTGGAAGCCACCAAGAGCCTGCTGGTGTGCAACAGCCGTATTCTCAGCTACATGGGTGAGCTGCAGCGGCGAAAGGAGGCCCAGGTGCTGGAGAAGACAG’GCGAGGGCAGGGGAAGCCTGCTCACCTGCGGCGACGTGGAGGAAAACCCTGGCCCCGCCCCAGGATCC |
| | **GFP-ROCK1 GBD** | |
| 27 | Xho1-F | TTCTCGAGCTCAAGATCAACTTGAAGCTGAG |
| 28 | BamH1-R | GCGGATCCCTAAGTTGAATCCGAAAGGTCCAAAAGTTTTGC |
| | **GFP-mDia1 GBD** | |
| 29 | Xho1-F | TTCTCGAGTAGAGCCGTCCGGCGGGGGCC |
| 30 | EcoR1-R | GCGAATTCCTACCCACTTTTTAATCCGTCCAGAAGTGGC |
