## Supplementary Materials for "Scaffolding of RhoA contractile signaling by anillin: a regulatory analogue of kinetic proofreading"

### Theoretical Supplement for "Scaffolding of RhoA contractile signalling by anillin: a regulatory analogue to kinetic proofreading"

This supplement describes calculations whose results are discussed in the main manuscript. Aside from providing sufficient detail for the results to be recapitulated, we also highlight important theoretical features of how an apparently inhibitory binding interaction between Anillin and RhoA can, in fact, promote downstream contractile signalling.

There are two main Sections: *"Anillin as a scaffold"* and *"Membrane phospholipids as a mechanism to antagonize dissociation"*. In the former, we describe the key facets of anillin's role as a scaffold, using a general semi-Markov representation. In the latter, we demonstrate how an explicit Markov reaction scheme, motivated in the main text, reproduces experimental data.

#### I. ANILLIN AS A SCAFFOLD

Consider a single GTP-RhoA molecule that has been recruited to the cortex. Once bound to the cortex, it may *i*) bind reversibly to a contractile effector (*e.g.*, ROCK1 or mDia1), *ii*) bind reversibly to anillin, or *iii*) dissociate from the cortex. Since anillin binds to the same domain of GTP-RhoA as effectors, it may *not* bind to GTP-RhoA while an effector is bound. Moreover, whilst GTP-RhoA is bound to either anillin or effectors, it is protected from cortical dissociation. The different states of the GTP-RhoA molecule, and the possible transitions between them are shown in Fig. 1**a**.

In the following, we first demonstrate how, in a simple Markovian realisation of the above single-molecule scheme, transient binding of anillin can increase the cortical residence time of GTP-RhoA, but only via sequestration: the number of interactions with effectors remains unchanged.

We then consider a generalisation of the same scheme, which is semi-Markov, having a dissociation step that is non-Poissonian. Here, transient binding is able to not only increase the total cortical residence time, but crucially, also the interaction with effectors.

##### A. Poissonian cortical dissociation

We first examine the case where all transitions have waiting-time distributions[1] that are exponential (*i.e.*, Poissonian), meaning that they can each be associated with a time-independent rate (see Fig. 1**b**). We use the labels $k_A$ ($d_A$) for binding (un-binding) with anillin, $k_E$ ($d_E$) for binding (un-binding) with effectors, and $k_D$ for cortical dissociation.

These single-molecule transition rates are related to the more common rate constants that appear in ordinary differential equation (ODE) treatments of mass-action kinetics. For example, the chemical reaction for the reversible binding of anillin to GTP-RhoA would usually be written as

$$\text{anillin } + \text{GTP-RhoA} \underset{k_b}{\overset{k_f}{\rightleftharpoons}} \text{RhoA-anillin Complex}, \tag{1}$$

where $k_f$ and $k_b$ are "forwards" and "backwards" rate constants. The corresponding mass-action ODE is just

$$\frac{d}{dt}[\text{RhoA-anillin Complex}] = k_f[\text{GTP-RhoA}][\text{anillin}] - k_b[\text{RhoA-anillin Complex}], \tag{2}$$

where square brackets $[\ldots]$ indicate concentration. Recall that the rate, $k_A$, of the single molecule transition $R \to A$ (*cf.* Fig. 1**a**) is just the rate that a given RhoA molecule meets any anillin molecule and forms a complex. Under the assumption that there is an sufficiently large external reservoir of anillin, we see that $k_A = k_f$ [anillin], which not only relates single molecule transition rates to mass-action rate constants, but also implies that $k_A \propto$ [anillin]. A similar argument can be used to show that $k_E \propto$ [effectors].

[1] For a transition $A \to B$, the waiting-time distribution $P_{A\to B}(t)$ is the distribution of probabilities associated with waiting a certain duration $t$, after arriving in state $A$, before moving to state $B$.

*1. Stabilization of GTP-RhoA by Anillin*

In the above context, the state of a single molecule is a continuous-time stochastic process, $X(t) \in \{R, A, E, D\}$, $\forall\, t > 0$. Writing $P_x(t) = \Pr\{X(t) = x\}$, the Master-Equation that governs the time evolution of the system can be represented as four coupled ODEs:

$$\begin{aligned}
\frac{\mathrm{d}P_A(t)}{\mathrm{d}t} &= k_A\, P_R(t) - d_A\, P_A(t),\\
\frac{\mathrm{d}P_E(t)}{\mathrm{d}t} &= k_E\, P_R(t) - d_E\, P_E(t),\\
\frac{\mathrm{d}P_R(t)}{\mathrm{d}t} &= k_A\, P_A(t) - d_A\, P_E(t) - (k_A + k_E + k_D)\; P_R(t),\\
\frac{\mathrm{d}\,P_D(t)}{\mathrm{d}t} &= k_D\, P_R(t),
\end{aligned} \tag{3}$$

with initial conditions $P_R(0) = 1$ and $P_A(0) = P_E(0) = P_D(0) = 0$. We wish to compute the mean cortical *residence* time— *i.e.*, the average time that a single molecule of GTP-RhoA takes to detach from the cortex following recruitment. This is nothing other than mean first passage time to dissociation. We write the distribution of such first passage times as

$$f_D(T) = \Pr\{X(T) = D,\ X(t) \neq D\ \forall\ 0 \leq t < T\}, \tag{4}$$

and therefore the mean is given by

$$\langle T_D \rangle = \int_0^\infty \mathrm{d}t\, t\, f_D(t). \tag{5}$$

The latter can be re-cast in terms of the Laplace transform:

$$\mathscr{L}\left[g(t)\right] = \bar{g}(s) = \int_0^\infty e^{-st}\, g(t)\, \mathrm{d}t, \tag{6}$$

where now

$$\langle T_D \rangle = -\left.\frac{\partial \overline{f_D}(s)}{\partial s}\right|_{s=0}. \tag{7}$$

We may also recognise that, since $D$ is an absorbing state, $f_D(t) = \mathrm{d}P_D(t)/\mathrm{d}t$, which implies

$$\langle T_D \rangle = -\left.\frac{\partial\left[s\,\overline{P_D}(s)\right]}{\partial s}\right|_{s=0}. \tag{8}$$

Laplace transforming Eqs. (3), and applying the aforementioned initial conditions, it is straightforward to show that

$$s\,\overline{P_D}(s) = \frac{k_D}{s - \left[\frac{k_A\, d_A}{s + d_A} + \frac{k_E\, d_E}{s + d_E} - (k_A + k_E + k_D)\right]}, \tag{9}$$

which can be substituted into (8) to obtain the result:

$$\langle T_D \rangle = \frac{1}{k_D}\left(1 + \frac{k_A}{d_A} + \frac{k_E}{d_E}\right). \tag{10}$$

That is, the mean cortical residence time increases with $k_A$, and hence the concentration of bulk anillin (Fig. 1**b**). This simply demonstrates that the more time GTP-RhoA spends bound to anillin, the more time is spent being blocked from cortical dissociation.

##### 2. *Anillin Stabilises GTP-RhoA by Sequestration*

We may also solve for the distribution of the number of reactions each GTP-RhoA molecule has with effectors before dissociation (*i.e.*, the number of times the state $E$ is visited in Fig. 1**a**). In order to do so, consider the sequence of states $\mathcal{S} = (R, x_1, x_2, \ldots, D)$ that are visited during any given realisation of the process $X(t)$. (Since there is an absorbing state, the process is guaranteed to terminate in finite time). The probability of a given sequence is just the sum of the probabilities of all trajectories $\mathcal{T} = ((R,0), (x_1,t_1), \ldots, (D,T))$ that correspond to the sequence $\mathcal{S}$. That is,

$$\Pr\{\mathcal{S} = (R, x_1, x_2, \ldots, D)\} = \int_{t_0<t_1<\ldots<t_n} \left(\prod_{i=1}^{n} \mathrm{d}t_i\right) \Pr\{\mathcal{T} = ((R,0), (x_1,t_1), \ldots, (D,T))\}. \tag{11}$$

The probability of a given trajectory can be written as a product of transition probabilities

$$\Pr\{\mathcal{T} = ((R,0), (x_1,t_1), \ldots, (D,T))\} = \prod_{i=0}^{n-1} Q_{x_i \to x_{i+1}}(t_{i+1} - t_i), \tag{12}$$

where $Q_{x_i \to x_j}$ is the probability that, given state $x_i$ at time $t_i$, the next transition is to state $x_j$ after time waiting for a period $t$— or more formally

$$\Pr\{X(t_i + t) = x_j, X(t_i + \tau) = x_i \;\forall\; \tau < t \mid X(t_i) = x_i\} = Q_{x_i \to x_j}(t). \tag{13}$$

Here, we exploit that the right-hand side of (11) is just a convolution over transition probabilities and hence, by the convolution theorem of Laplace transforms

$$\Pr\{\mathcal{S} = (R, x_1, x_2, \ldots, D)\} = \prod_{i=0}^{n-1} \overline{Q}_{x_i \to x_j}(0), \tag{14}$$

where from (6)

$$\overline{Q}_{x_i \to x_j}(0) = \int_0^\infty Q_{x_i \to x_j}(t)\,\mathrm{d}t. \tag{15}$$

Going back to (13), it is clear that $Q_{x_i \to x_j}(t)$ further factors into the product of a waiting time distribution

$$\Pr\{X(t_i + t) = x_j \mid X(t_i + \tau) = x_i \;\forall\; \tau < t,\; X(t_i) = x_i\} = P_{x_i \to x_j}(t), \tag{16}$$

and the probability that nothing else occurs during for the duration of the waiting time

$$\Pr\{X(t_i + \tau) = x_i \;\forall\; \tau < t, \mid X(t_i) = x_i\} = \prod_{x_k \neq x_i, x_j} S_{x_i \to x_k}(t), \tag{17}$$

where

$$S_{x_i \to x_k}(t) = 1 - \int_0^t \mathrm{d}\tau\, P_{x_i \to x_j}(\tau), \tag{18}$$

is a survivor function. We may now substitute for the known (Poissonian) waiting time distributions: $P_{R\to x} = k_x\, e^{-k_x\, t},\ \forall x \in \{A, E, D\}$ and $P_{x \to R} = d_x\, e^{-d_x\, t},\ \forall x \in \{A, E\}$, to obtain

$$\overline{Q}_{R\to x}(0) = \frac{k_x}{k_A + k_E + k_D},\ \forall\; x \in \{A, E, D\},\ \text{and}\ \overline{Q}_{x\to R}(0) = 1,\ \forall\; x \in \{A, E\}. \tag{19}$$

Using the integers $m$ and $l$, we write

$$\Omega_{m,l} = \left\{\mathcal{S} = (R, x_1, \ldots, D) : \{x_1, \ldots, x_{n-1}\} = \left\{E^m, A^l, R^{n-(m+l+1)}\right\}\right\}, \tag{20}$$

for the set of all sequences under which the state $E$ is visited $m$ times and the state $A$ is visited $l$ times. The probability of realising a sequence belonging to $\Omega_{m,l}$ is then

$$\Pr\{\mathcal{S} = \omega \in \Omega_{m,l}\} = \left[\overline{Q}_{R\to E}(0)\right]^m \left[\overline{Q}_{R\to A}(0)\right]^l \left[\overline{Q}_{R\to D}(0)\right]. \tag{21}$$

Combinatorially accounting for the ordering of transitions, it is then clear that the probability of visiting the state $E$ on $m$ distinct occasions is just,

$$\begin{aligned}\Pr\{\mathcal{S}=\lambda\in\Lambda_m\} &= \sum_l \binom{l+m}{l} \left[\overline{Q}_{R\to E}(0)\right]^m \left[\overline{Q}_{R\to A}(0)\right]^l \left[\overline{Q}_{R\to D}(0)\right] \\ &= \frac{k_D}{k_E+k_D}\left(\frac{k_E}{k_E+k_D}\right)^m,\end{aligned} \tag{22}$$

where we use

$$\Lambda_m = \left\{\mathcal{S}=(R,x_1,\ldots,D):\{x_1,\ldots,x_{n-1}\}=\left\{E^m,A^l,R^{n-(m+l+1)}\right\}\ \forall\ l\in\mathbb{N}\right\}. \tag{23}$$

to denote the set of all sequences under which the state $E$ is visited $m$ times. Importantly, this result does not depend on $k_A$, the rate of complex formation with anillin. As a result, the mean number of interactions with effectors:

$$\langle m\rangle = \sum_m \frac{m\,k_D}{k_E+k_D}\left(\frac{k_E}{k_E+k_D}\right)^m = \frac{k_E}{k_D}, \tag{24}$$

is also independent of $k_A$. Since the mean residence time increases with $k_A$ whilst the mean number of interactions with effectors does not, the mean number of interactions per molecule per unit time *decreases* with $k_A$. The above procedure can also be used to calculate the average number of visits to the state $A$:

$$\begin{aligned}\Pr\{\mathcal{S}=\lambda\in\Lambda_l\} &= \sum_m \binom{l+m}{m} \left[\overline{Q}_{R\to E}(0)\right]^m \left[\overline{Q}_{R\to A}(0)\right]^l \left[\overline{Q}_{R\to D}(0)\right] \\ &= \frac{k_D}{k_A+k_D}\left(\frac{k_A}{k_A+k_D}\right)^l,\end{aligned} \tag{25}$$

where

$$\Lambda_l = \left\{\mathcal{S}=(R,x_1,\ldots,D):\{x_1,\ldots,x_{n-1}\}=\left\{E^m,A^l,R^{n-(m+l+1)}\right\}\ \forall\ m\in\mathbb{N}\right\}, \tag{26}$$

which implies

$$\langle l\rangle = \frac{k_A}{k_D}. \tag{27}$$

The average time spent in the anillin bound state, $\langle\tau_A\rangle$, is then just $\langle l\rangle$ multiplied by the average time spent there: $1/d_A$. The result is that

$$\langle\tau_A\rangle = \frac{k_A}{k_D\,d_A}, \tag{28}$$

which demonstrates that any increase in the cortical residence time of GTP-RhoA due to anillin is solely due to an increase in the mean time spent in the anillin bound state (Fig. 1**c**). That is, in this model, anillin stabilises RhoA by *sequestration*, blocking it not only from cortical dissociation, but also interactions with effectors.

##### 3. *Increasing Anillin Concentration Does Not Affect Signalling to Effectors*

For both completeness and in order to compare with experimental measurements, we consider a system with many GTP-RhoA molecules, able to associate and dissociate to- and from- the cortex, respectively. When bound, each molecule is subject to the Poissonian version of the reaction scheme in Fig. 1**a**. We assume the molecules are non-interacting, neither directly nor indirectly. For the latter, we are required to assume that the bulk reservoirs of anillin, effectors, and GDI are all sufficiently large that their concentrations, and hence the rates $k_A$, $k_E$, and $k_D$, are unchanged by what goes on at the cortex.

The number of molecules at the cortex is now a stochastic variable $N(t)$. We invoke a standard result from queuing theory that, given a Poissonian cortical recruitment process with rate $k_{\text{on}}$, and a mean first passage time to dissociation $\langle T_D\rangle$ [*i.e.*, Eq. (10)], then the distribution of the number of molecules $n$ at the cortex is a Poisson distribution:

$$\Pr\{N(t)=n\} = \frac{\left(k_{\text{on}}\langle T_D\rangle\right)^n e^{-\left(k_{\text{on}}\langle T_D\rangle\right)}}{n!}. \tag{29}$$

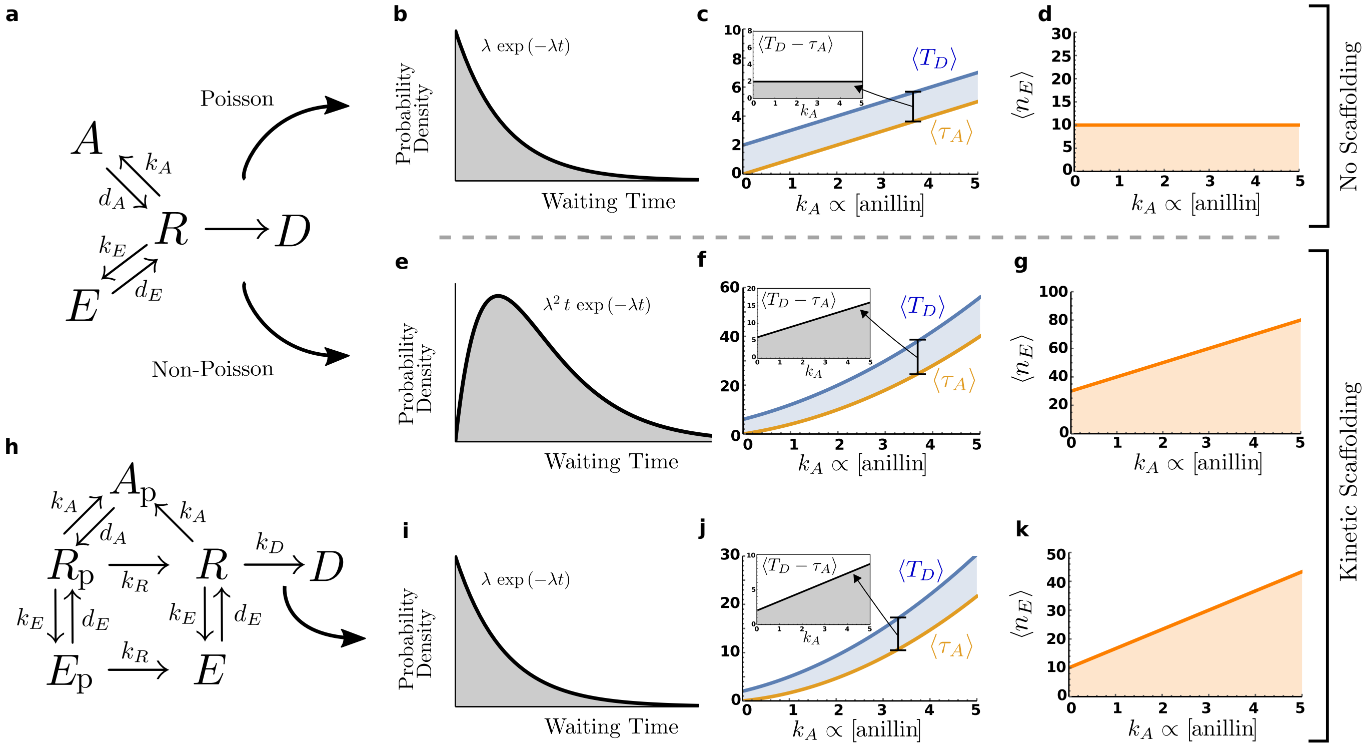


FIG. 1. **Anillin as a scaffold**. **a**) The simplest single molecule reaction scheme for GTP-RhoA which, in its free state at the cortex ($R$) can bind to anillin ($A$), to effectors ($E$), and dissociate from the membrane ($D$). The labels $k_x$ and $d_x$ represent rates for transitions into and out of states $x \in \{A, E\}$. **b**) A generic (Poissonian) exponential waiting time distribution, characterised by rate $\lambda$. **c**) The mean residence time of a single GTP-RhoA molecule, $\langle T_D \rangle$, and the mean time spent bound to anillin, $\langle \tau_A \rangle$, as a function of anillin concentration. The inset shows the difference, demonstrating that any gain is residence time due to sequestration by anillin. **d**) Correspondingly, the mean number of effector bound GTP-RhoA molecules at the cortex, $\langle n_E \rangle$, is independent of anillin concentration. **e**) A representative non-Poissonian waiting time distribution for dissociation $R \to D$, here chosen to be an Erlang distribution with shape parameter $n = 2$. **f**) The residence time $\langle T_D \rangle$ and sequestered time $\langle \tau_A \rangle$ for the dissociation process as shown in **e**. In contrast to **b**, the difference $\langle T_D - \tau_A \rangle$ grows with anillin concentration. **g**) Correspondingly, the number of effector bound GTP-RhoA molecules also increases with anillin concentration. **h**) The single molecule network incorporating interactions with PIP2 (see main text). In contrast to panels **e** through **g**, all transitions are Poissonian, with an exponential waiting time distribution as shown in **i**. **j**) The residence time $\langle T_D \rangle$ and sequestered time $\langle \tau_A \rangle$ once again increase with anillin concentration; as does the difference $\langle T_D - \tau_A \rangle$ (inset). **k**) the number of effector bound GTP-RhoA molecules (*i.e.*, those in either of states $E$ or $E_\mathrm{p}$) also increases with anillin concentration.

Moreover, the average number of effector-bound molecules, $\langle n_E \rangle$, is just the average number of molecules, $\langle n \rangle = k_\mathrm{on} \langle T_D \rangle$, multiplied by the fraction of time each spends in state $E$ on average, $\langle \tau_E \rangle / \langle T_D \rangle$. The result is that

$$\langle n_E \rangle = \frac{k_E \, k_\mathrm{on}}{k_D \, d_E}, \tag{30}$$

which is independent of $k_A$, and hence the concentration of bulk anillin (Fig. 1**d**).

##### B. *Non*-Poissonian Cortical Dissociation

The results of the previous Section are at odds with the experimental evidence that, despite sharing a RhoA binding domain with contractile effectors, direct binding of anillin can simultaneously stabilize RhoA at the cortex and promote contractile signalling. To resolve this apparent paradox, we are led to propose a *semi*-Markov approach, whereby the waiting-time distribution for the dissociation step, $P_{R \to D}(t)$, is no longer exponential, or *non*-Poissonian (Fig. 1**e**).

##### 1. Enhanced Stabilization of GTP-RhoA by Anillin

To compute the first passage time to dissociation, we can no longer use the Master-Equation. Instead, we adapt the path summation method used in Sec. I A 2. Here, the relevant quantity is not the probability that a given realisation of $X(t)$ results in a particular sequence of states, $\mathcal{S}$. Rather, we want the joint probability of given sequence and its dissociation time $T$. Using the same notation as the previous section, we write

$$\Pr\left\{\mathcal{S}=(R,x_1,\ldots,D),t_n=T\right\}=\int_{t_0<t_1<\ldots<t_{n-1}}\left(\prod_{i=1}^{n-1}\mathrm{d}t_i\right)\prod_{i=0}^{n-2}Q_{x_i\to x_{i+1}}(t_{i+1}-t_i)\,Q_{x_{n-1}\to D}(t_{n-1}-T). \tag{31}$$

where, in contrast to (11), there is no integration over the final time to dissociation. This can be expressed in more compact form by applying the Laplace transform and invoking the convolution theorem:

$$\mathscr{L}\left[\Pr\left\{\mathcal{S}=(R,x_1,\ldots,D),t_n=T\right\}\right]=\prod_{i=0}^{n-1}\overline{Q}_{x_i\to x_{i+1}}(s), \tag{32}$$

where the variable $s$ is conjugate to the dissociation time $T$. As before, the probability of a certain sequence only depends on how many times the states $E$ and $A$ were visited. Using (20), we have

$$\mathscr{L}\left[\Pr\left\{\mathcal{S}=\omega\in\Omega_{m,l},t_n=T\right\}\right]=\left[\overline{Q}_{R\to E}(s)\,\overline{Q}_{E\to R}(s)\right]^m\,\left[\overline{Q}_{R\to A}(s)\,\overline{Q}_{A\to R}(s)\right]^l\,\overline{Q}_{R\to D}(s). \tag{33}$$

where the Laplace transform of the distribution of first passage times can be obtained by marginalising over all sequences:

$$\begin{aligned}\overline{f}_D(s)&=\sum_{m,l}\binom{l+m}{m}\,\left[\overline{Q}_{R\to E}(s)\,\overline{Q}_{E\to R}(s)\right]^m\,\left[\overline{Q}_{R\to A}(s)\,\overline{Q}_{A\to R}(s)\right]^l\,\overline{Q}_{R\to D}(s)\\&=\frac{\overline{Q}_{R\to D}(s)}{1-\left[\overline{Q}_{R\to E}(s)\,\overline{Q}_{E\to R}(s)+\overline{Q}_{R\to A}(s)\,\overline{Q}_{A\to R}(s)\right]}.\end{aligned} \tag{34}$$

Exploiting (7), the mean first passage time is then

$$\langle T_D\rangle=-\frac{\partial}{\partial s}\left\{\frac{\overline{Q}_{R\to D}(s)}{1-\left[\overline{Q}_{R\to E}(s)\,\overline{Q}_{E\to R}(s)+\overline{Q}_{R\to A}(s)\,\overline{Q}_{A\to R}(s)\right]}\right\}\Bigg|_{s=0}. \tag{35}$$

As before, $P_{R\to x}=k_x\,e^{-k_x\,t}$, $\forall\ x\in\{A,E\}$, and $P_{x\to R}=d_x\,e^{-d_x\,t}$, $\forall\ x\in\{A,E\}$. However, the waiting-time distribution associated with the transition $R\to D$ can now be arbitrarily chosen. We choose an Erlang distribution with general shape parameter $n$

$$P_{R\to D}=\frac{k_D^n\,t^{(n-1)}\,e^{-k_D\,t}}{(n-1)!}, \tag{36}$$

which corresponds to a chain of $n$ Poisson processes, (*i.e.*, $n-1$ intermediates). The result is that

$$\overline{Q}_{R\to x}(s)=\frac{k_x}{s+k_A+k_E}\left[1-\left(\frac{k_D}{s+k_A+k_E+k_D}\right)^n\right],\ \forall\ x\in\{A,E\}, \tag{37}$$

$$\overline{Q}_{x\to R}(s)=\frac{d_x}{s+d_x},\ \forall\ x\in\{A,E\}, \tag{38}$$

and

$$\overline{Q}_{R\to D}(s)=\left(\frac{k_D}{s+k_A+k_E+k_D}\right)^n. \tag{39}$$

Substituting into (35), we see that the mean first passage time to dissociation is then

$$\langle T_D\rangle=\frac{\left[d_E\,k_A+d_A\,(d_E+k_E)\right]}{d_A\,d_E\,(k_A+k_E)}\left[\left(\frac{k_A+k_D+k_E}{k_D}\right)^n-1\right], \tag{40}$$

which increases $\sim k_A^n$. That is, for the case of non-Poissonian dissociation ($n\geq 2$) the cortical residence time not only increases with bulk anillin concentration, but the effect is enhanced over that of the simple Poissonian case. This is shown explicitly for the case of a single intermediate inactivation step (*i.e.*, $n=2$) in Fig. 1**f**.

*2. Anillin Increases the Dwell Time of "Free" GTP-RhoA*

To understand whether the increase in $\langle T_D \rangle$ is simply due to sequestration (as in the case of Poissonian dissociation), we calculate $\langle m \rangle$ and $\langle l \rangle$, the mean number of interactions with effectors and anillin, respectively. For the former, we may substitute for the expressions (37-39) in (22) and take the average over $m$, with the result

$$\begin{aligned}\langle m \rangle &= \sum_{l,m} \binom{l+m}{l} m \left[\overline{Q}_{R\to E}(0)\right]^m \left[\overline{Q}_{R\to A}(0)\right]^l \left[\overline{Q}_{R\to D}(0)\right] \\ &= \frac{k_E}{k_A + k_E}\left[\left(\frac{k_A + k_D + k_E}{k_D}\right)^n - 1\right].\end{aligned} \tag{41}$$

Similarly, for the latter

$$\begin{aligned}\langle l \rangle &= \sum_{m,l} \binom{l+m}{m} l \left[\overline{Q}_{R\to E}(0)\right]^m \left[\overline{Q}_{R\to A}(0)\right]^l \left[\overline{Q}_{R\to D}(0)\right] \\ &= \frac{k_A}{k_A + k_E}\left[\left(\frac{k_A + k_D + k_E}{k_D}\right)^n - 1\right].\end{aligned} \tag{42}$$

Equation (41) indicates that as $k_A$ increases, so does the average number of interactions a single GTP-RhoA has with effectors. Equation (42) can be used to calculate the average time spent in the anillin bound state, $\langle \tau_A \rangle = \langle l \rangle / d_A$, and hence the increase in residence time *not* associated with sequestration by anillin:

$$\langle T_D - \tau_A \rangle = \frac{d_E + k_E}{d_E\,(k_A + k_E)}\left[\left(\frac{k_A + k_E + k_D}{k_D}\right)^n - 1\right]. \tag{43}$$

For $n \geq 2$, the above expression $\sim k_A^n$ (see Fig. 1**f**).

*3. Increasing Anillin Concentration Promotes Signalling to Effectors*

Once again moving to a picture of the cortex, we imagine a large number of the aforementioned semi-Markov processes occurring in parallel, where binding to the cortex from the bulk is a simple Poisson process. Using (29), we can calculate the average number of GTP-RhoA molecules that are in an effector-bound state at any given time:

$$\langle n_E \rangle = k_{\rm on}\langle \tau_E \rangle = \frac{k_{\rm on}\,\langle m \rangle}{d_E} = \frac{k_{\rm on}\,k_E}{d_E\,(k_A + k_E)}\left[\left(\frac{k_A + k_D + k_E}{k_D}\right)^n - 1\right], \tag{44}$$

from which it is clear that signalling of Rho to effectors (at constant bulk concentration of RhoA) can be increased by increasing the bulk concentration of anillin, and hence $k_A$ (Fig. 1**g**).

#### II. MEMBRANE PHOSPHOLIPIDS AS A MECHANISM TO ANTAGONISE DISSOCIATION

The previous section demonstrated that the experimental observations could be explained by assuming a very simple single-molecule reaction network and invoking a non-Poissonian process of cortical dissociation. Such non-Poissonian behaviour is likely a coarse-grained representation of underlying intermediate states and transitions which, for resetting-like behaviour, form at least one closed loop, or cycle, in state-space. The experiments described in the main manuscript suggested that the localisation of acidic phospholipids by anillin might lead to such intermediate steps. Therefore, this section describes an explicit Markov reaction scheme, involving PIP2 as a representative acidic phospholipid, which exhibits the same features of stochastic resetting as the semi-Markov processes of Sec. I B.

Here, we recognise our experimental observation that clustering the AH domain (IL2R-AH) locally co-accumulated PIP2 and note that acidic phospholipids can antagonise cortical dissociation of RhoA, both directly and indirectly.

Our approach assumes a dilute background of PIP2, with a local enhancement of concentration around anillin, such that the probability of a free GTP-RhoA forming a complex with PIP2 unaided is negligible (*i.e.*, the reaction $R \to R_{\rm p}$ is excluded). If a free GTP-RhoA binds to anillin, however, then it is temporarily held in an area of high local PIP2 concentration, implying that, on unbinding from anillin, GTP-RhoA has a high probability of being bound to PIP2: (*i.e.*, $R \to A_{\rm p}$ followed by $A_{\rm p} \to R_{\rm p}$, with $A_{\rm p} \to R$ disallowed). The transient binding of GTP-RhoA to PIP2 provides temporary protection from dissociation (*i.e.*, $R_{\rm p} \to D$ is not allowed).

The relevant reaction scheme is set out in Fig. 1**h**. Here, we may use much of the machinery introduced in the previous sections, however, due to the presence of a cycle, the use of a simple Binomial coefficient is not sufficient for counting the contributions from possible paths. We therefore introduce the matrix $\mathsf{A}$, whose coefficients are defined by

$$A_{ij} = \overline{Q}_{y_i \to y_j}(s) = \begin{pmatrix} 0 & \frac{k_E}{k_A+k_E+k_D+s} & \frac{k_A}{k_A+k_E+k_D+s} & 0 & 0 & \frac{k_D}{k_A+k_E+k_D+s} \\ \frac{d_E}{d_E+s} & 0 & 0 & 0 & 0 & 0 \\ 0 & 0 & 0 & \frac{d_A}{d_A+s} & 0 & 0 \\ \frac{k_R}{k_A+k_E+k_R+s} & 0 & \frac{k_A}{k_A+k_E+k_R+s} & 0 & \frac{k_E}{k_A+k_E+k_R+s} & 0 \\ 0 & \frac{k_R}{d_E+k_R+s} & 0 & \frac{d_E}{d_E+k_R+s} & 0 & 0 \\ 0 & 0 & 0 & 0 & 0 & 0 \end{pmatrix}, \tag{45}$$

where the labelling convention is that: $y_1 = R$, $y_2 = E$, $y_3 = A_\mathrm{p}$, $y_4 = R_\mathrm{p}$, $y_5 = E_\mathrm{p}$, $y_6 = D$. The $i = 1$, $j = 6$ entry of $\mathsf{A}$ is just the Laplace transform of the distribution of first passage times to dissociation *made with only one transition.* The $i = 1$, $j = 6$ entry of $\mathsf{A}^2$ is just the Laplace transform of the distribution of first passage times to dissociation *made with only two transitions.* The $i = 1$, $j = 6$ entry of $\mathsf{A}^n$ is just the Laplace transform of the distribution of first passage times to dissociation *made with only $n$ transitions.* Therefore taking a so-called Neumann-series, it can be seen that

$$\overline{f}_D(s) = \left[\sum_{n=0}^{\infty} \mathsf{A}^n\right]_{i=1,j=6} = \left[(\mathbb{I} - \mathsf{A})^{-1}\right]_{i=1,j=6}, \tag{46}$$

where the subscript indicates taking the $(1, 6)$ component of the resulting matrix. Using (7), it then follows that

$$\langle T_D \rangle = \frac{[d_E\, k_A + d_A\,(d_E + k_E)]\,[d_E\,(k_A + k_R) + k_R\,(k_A + k_E + k_R)]}{d_A\, d_E\, k_D\, k_R\,(d_E + k_E + k_R)}. \tag{47}$$

To calculate the quantity $\langle m \rangle$, we introduce the matrix $\mathsf{B}(z)$ such that

$$B_{ij}(z) = \begin{cases} z\, A_{ij}(0) & \forall\ (i,j) \in \{(1,2),(4,5)\} \\ A_{ij}(0) & \text{otherwise} \end{cases}. \tag{48}$$

That is, the variable $z$ appears in each entry corresponding to a transition into either state $E_\mathrm{p}$ or state $E$. The corresponding Neumann-series turns out to be the generating function for the distribution $P(m) = \Pr\{\mathcal{S} = \lambda \in \Lambda_m\}$. That is

$$G(z) = \sum_{m=0}^{\infty} z^m\, P(m) = \left[\sum_{n=0}^{\infty} \mathsf{B}^n\right]_{i=1,j=6} = \left[(\mathbb{I} - \mathsf{B})^{-1}\right]_{i=1,j=6}. \tag{49}$$

As a result, it follows that

$$\langle m \rangle = \left.\frac{\partial G}{\partial z}\right|_{z=1} = \frac{k_E\,[d_E\,(k_A + k_R) + k_R\,(k_A + k_E + k_R)]}{k_D\, k_R\,(d_E + k_E + k_R)}, \tag{50}$$

which increases with $k_A \propto$ [anillin], as expected. In a similar way, we may define $\mathsf{C}$ such that

$$C_{ij}(z) = \begin{cases} z\, A_{ij}(0) & \forall\ (i,j) \in \{(1,3),(4,3)\} \\ A_{ij}(0) & \text{otherwise} \end{cases}, \tag{51}$$

where the variable $z$ now appears in each entry corresponding to a transition into $A_\mathrm{p}$. As a result

$$H(z) = \sum_{m=0}^{\infty} z^l\, P(l) = \left[\sum_{n=0}^{\infty} \mathsf{C}^n\right]_{i=1,j=6} = \left[(\mathbb{I} - \mathsf{C})^{-1}\right]_{i=1,j=6}. \tag{52}$$

is now the generating function for $P(l) = \Pr\{\mathcal{S} = \lambda \in \Lambda_l\}$, where

$$\langle l \rangle = \left.\frac{\partial H}{\partial z}\right|_{z=1} = \frac{k_A\,[d_E\,(k_A + k_R) + k_R\,(k_A + k_E + k_R)]}{k_D\, k_R\,(d_E + k_E + k_R)}. \tag{53}$$

Substituting into $\langle \tau_A \rangle = \langle l \rangle / d_A$ and using (47), it can be shown that the mean time spent at the cortex not in the anillin-bound state

$$\langle T_D - \tau_A \rangle = \frac{(d_E + k_E)\left[d_E\,(k_A + k_R) + k_R\,(k_A + k_E + k_R)\right]}{d_E\,k_D\,k_R\,(d_E + k_E + k_R)}. \tag{54}$$

also increases with $k_A \propto$ [anillin] (Fig. 1**j**). Moreover, using (50), we see that, in the context of the cortical picture described earlier

$$\langle n_E \rangle = \frac{k_{\rm on}\,\langle m \rangle}{d_E} = \frac{k_{\rm on}\,k_E}{d_E\,(k_A + k_D)}\left[\left(\frac{k_A + k_D + k_E}{k_D}\right)^n - 1\right]. \tag{55}$$

That is, the mean number of GTP-RhoA molecules in the effector-bound state (at any given time) increases with $k_A \propto$ [anillin], as expected (Fig. 1**k**).

Here, we highlight the fact that, by introducing a simple cycle, our network resembles the Hopfield-Ninio proofreading motif. There, a cycle implies a mean first-passage time— for "activating" a given cellular response— that is quadratic in the dissociation rate of a particular enzyme-substrate pair, implying greater discriminatory, or "proofreading", ability. [For example, low-affinity substrates (with high dissociation rates) lead to especially low rates of "activation"]. In our "regulatory" case, the analogous effect is the quadratic dependence of the mean residence time on the rate, $k_A$, with which anillin binds to GTP-RhoA. Our schema seeks to exploit a corollary of this fact; that the mean time spent *unbound* to anillin is linear in $k_A$ (rather than independent, as would be the case with no cycles).
